# Learning compositional sequences with multiple time scales through a hierarchical network of spiking neurons

**DOI:** 10.1101/2020.09.08.287748

**Authors:** Amadeus Maes, Mauricio Barahona, Claudia Clopath

## Abstract

Sequential behaviour is often compositional and organised across multiple time scales: a set of individual elements developing on short time scales (motifs) are combined to form longer functional sequences (syntax). Such organisation leads to a natural hierarchy that can be used advantageously for learning, since the motifs and the syntax can be acquired independently. Despite mounting experimental evidence for hierarchical structures in neuroscience, models for temporal learning based on neuronal networks have mostly focused on serial methods. Here, we introduce a network model of spiking neurons with a hierarchical organisation aimed at sequence learning on multiple time scales. Using biophysically motivated neuron dynamics and local plasticity rules, the model can learn motifs and syntax independently. Furthermore, the model can relearn sequences efficiently and store multiple sequences. Compared to serial learning, the hierarchical model displays faster learning, more flexible relearning, increased capacity, and higher robustness to perturbations. The hierarchical model redistributes the variability: it achieves high motif fidelity at the cost of higher variability in the between-motif timings.

## INTRODUCTION

Many natural behaviours are compositional: complex patterns are built out of combinations of a discrete set of simple motifs (Tresch et al. 1999; Bizzi et al. 2008; Wiltschko et al. 2015). Compositional sequences unfolding over time naturally lead to the presence of multiple time scales—a short time scale is associated with the motifs and a longer time scale is related to the ordering of the motifs into a syntax. How such behaviours are learnt and controlled is the focus of much current research. Broadly, there are two main strategies for the modeling of sequential behaviour: serial and hierarchical. In a serial model, the long-term behaviour is viewed as a chain of motifs proceeding sequentially, so that the first behaviour in the chain leads to the second and so on (‘domino effect’). Serial models present some limitations (Lashley 1951; Houghton and Hartley 1995). Firstly, serial models have limited flexibility since relearning the syntax involves rewiring the chain. Secondly, such models lack robustness, e.g., breaking the serial chain halfway means that the later half of the behaviour is not produced. It has been proposed theoretically that hierarchical models can alleviate these problems, at the cost of extra hardware.

Evidence for the presence of hierarchical structures in the brain is mounting (Tanji 2001; Kiebel et al. 2008; Murray et al. 2014). Furthermore, experiments are increasingly shining light on the hierarchical mechanisms of sequential behaviour. An example is movement sequences in multiple animal models, such as *Drosophila* (Seeds et al. 2014; Berman et al. 2016; Jovanic et al. 2016), mice (Jin and Costa 2015; Geddes et al. 2018; Markowitz et al. 2018) and *C. elegans* (Kato et al. 2015; Kaplan et al. 2020). Simultaneous recordings of behaviour and neural activity are now possible in order to relate the two together (Vogelstein et al. 2014; Berman 2018). Songbirds are another example of animals that produce stereotypical sequential behaviour: short motifs are strung together to form songs. In this case, a clock-like dynamics is generated in the premotor nucleus HVC of the bird’s brain, such that neurons are active in sequential bursts of ∼ 10 ms (Hahnloser et al. 2002). This activity is thought to control the timing of the spectral content of the song (the within-motif dynamics). The between-motif dynamics has a different temporal structure (Glaze and Troyer 2006; Glaze and Troyer 2013); hence the ordering of the motifs into a song (the syntax) might be controlled by a different mechanism. Supporting this view, it has been found that learning the motifs and syntax involves independent mechanisms (Lipkind et al. 2017). The computational study of hierarchical structures and compositional behaviour can also lead to insights into the development of human locomotion and language as there are striking conceptual parallels (Dominici et al. 2011; Lipkind et al. 2013; Ding et al. 2015; Lipkind et al. 2019).

Here, we present a model for learning temporal sequences on multiple scales implemented through a hierarchical network of bio-realistic spiking neurons and synapses. In contrast to current models, which focus on acquiring the motifs and speculate on the mechanisms to learn a syntax (Stroud et al. 2018; Logiaco et al. 2019; Maes et al. 2020), our spiking network model learns motifs and syntax independently from a target sequence presented repeatedly. Furthermore, the plasticity of the synapses is entirely local, and does not rely on a global optimisation such as FORCE-training (Nicola and Clopath 2017; Hardy and Buonomano 2018; Nicola and Clopath 2019) or backpropagation through time (Werbos 1990). To characterise the effect of the hierarchical organisation, we compare the proposed hierarchical model to a serial version by looking at their learning and relearning behaviours. We show that, contrary to the serial model, the hierarchical model acquires the motifs independently from the syntax. In addition, the hierarchical model has a higher capacity and is more resistant to perturbations, as compared to a serial model. We also investigate the variability of the neural activity in both models, during spontaneous replay of stored sequences. The organisation of the model shapes the neural variability differently. The within-motif spiking dynamics is less variable in a hierarchical organisation, while the time between the execution of motifs is more variable.

The paper is organised as follows. We start by describing the proposed hierarchical spiking network model and the learning protocol. We then analyse the learning and relearning behaviour of the proposed model, and compare it to the corresponding serial model. Next, we investigate several properties of the model: (i) the performance and consistency of spontaneous sequence replays on a range of learnt sequences; (ii) capacity, i.e., how multiple sequences can be stored simultaneously; (iii) robustness of the sequence replays.

## RESULTS

### Hierarchical model of spiking neurons with plastic synapses for temporal sequence learning

We design a hierarchical model by combining the following spiking recurrent networks (Fig. 1): 1) A recurrent network exhibiting fast sequential dynamics (the *fast clock*); 2) a recurrent network exhibiting slow sequential dynamics (the *slow clock*); 3) a series of interneuron networks that store and produce the to-be-learnt ordering of motifs (the *syntax networks*); 4) a series of read-out networks that store and produce the to-be-learnt motif dynamics (the *motif networks*). We assume that there are a finite number of motifs and each motif is associated to a separate read-out network (e.g., in Fig. 1 there are 2 read-out networks corresponding to motifs *A* and *B*). The goal of the model is to learn a complex sequence, with the motifs arranged in a certain temporal order, such that the motifs themselves and the temporal ordering of the motifs are learnt using local plasticity rules.

**Fig. 1.**
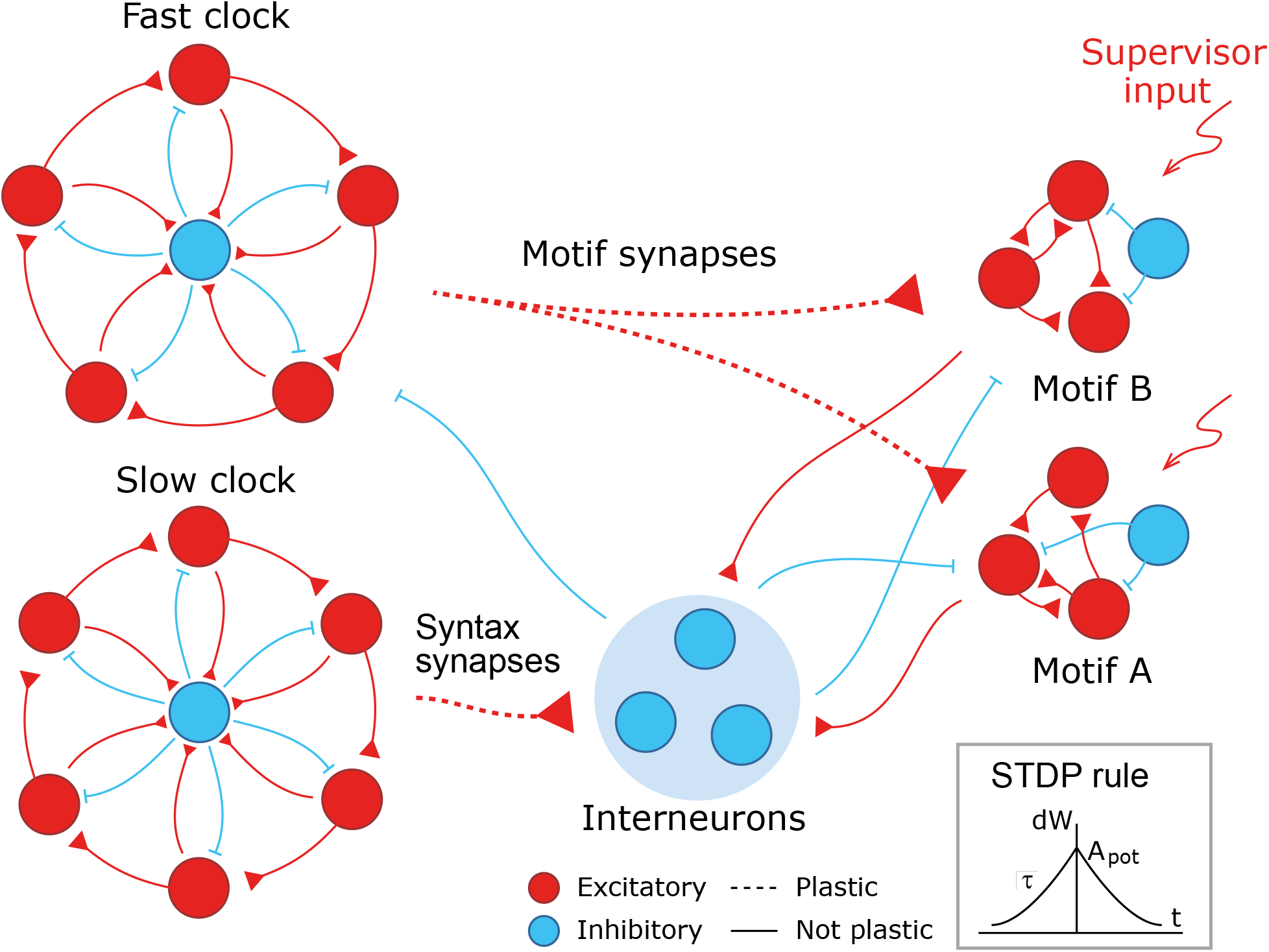
A cartoon of the model. Dynamics in the read-out networks (A and B) is learnt and controlled on two time scales. The fast time scale network (fast clock) exhibits sequential dynamics that spans individual motifs. This acts directly on the read-out networks through plastic synapses. These synapses learn the *motifs*. The slow time scale network (slow clock) exhibits sequential dynamics that spans the entire sequence of motifs. This acts indirectly on the read-out networks through an interneuron network. The synapses from the slow clock to the interneurons are plastic and learn the right order of the motifs, or the *syntax*. The plastic synapses follow a simple symmetric STDP rule for potentiation, with a constant depression independent of spike time.

#### Neuronal network architecture

All neurons are either excitatory or inhibitory. Excitatory neurons follow an adaptive exponential integrate-and-fire dynamics and inhibitory neurons follow a standard integrate-and-fire dynamics (see Methods).

The model has two recurrent networks that exhibit sequential dynamics: the fast and slow clocks. The design of the clock networks follows Ref. Maes et al. 2020. Each clock is composed of clusters of excitatory neurons coupled in a cycle with a directional bias (i.e., neurons in cluster *I* are more strongly connected to neurons in cluster *i* + 1) together with a central cluster of inhibitory neurons coupled to all the excitatory clusters (Fig. 1). This architecture leads to sequential dynamics propagating around the cycle and the period can be tuned by choosing different coupling weights. The individual motifs are not longer than the period of the fast clock and the total length of the sequence is limited to the period of the slow clock. In our case, we set the coupling weights of the fast clock such that a period of ∼ 200 ms is obtained, whereas the weights of the slow clock are set to obtain a period of ∼ 1000 ms.

The fast clock neurons project directly onto the read-out networks associated with each motif, which are learnt and encoded using a supervisor input. Hence the fast clock controls the within-motif dynamics. The slow clock neurons, on the other hand, project onto the interneuron network of inhibitory neurons. The interneuron network is also composed of clusters: there is a cluster associated with each motif, with coupling weights that inhibit all other motif networks, and one cluster associated with the ‘silent’ motif, with couplings that inhibit all motif networks and the fast clock. Hence the temporal ordering of the motifs (the syntax) can be encoded in the mapping that controls the activity of the interneurons driven by the slow clock. As a result of this hierarchical architecture, the model allows for a dissociation of within-motif dynamics and motif ordering. The two pathways, from the fast clock to the read-out and from the slow clock to the interneurons, each control a different time scale of the spiking network dynamics.

#### Plasticity

Learning is accomplished through plastic synapses under a simple biophysically plausible local STDP rule (see Methods) governing the synapses from the fast clock to the read-out networks (motif synapses) and from the slow clock to the interneurons (syntax synapses). The STDP rule has a symmetric learning window and implements a Hebbian ‘fire together, wire together’ mechanism.

All other weights in the model are not plastic and are fixed prior to the learning protocol. The weights in the fast and slow clocks and the interneuron wiring are assumed to originate from earlier processes during evolution or early development. Previous computational studies have shown that sequential dynamics can be learnt in recurrent networks, both in an unsupervised (Jun and Jin 2007; Zheng and Triesch 2014) and supervised (Murray and Escola 2017; Maes et al. 2020) fashion.

#### Learning scheme

During learning, a target sequence is presented. We design a target sequence by combining motifs in any order, e.g., *AAB*. A time-varying external current, corresponding to the target sequence, projects to the excitatory neurons in the read-out networks. Additionally, a short external current activates the first cluster in the fast clock to signal the onset of a new motif (see Methods for more details). During the presentation of the target sequence, the plastic synapses change. When no target sequence is presented, spontaneous dynamics is simulated. Spontaneous dynamics replays the stored sequence. In this case, there is only random external input and no external input corresponding to a target sequence.

### The model allows for independent learning of motifs and syntax

We first show how a non-trivial sequence can be learned emphasising the role that each network plays. As an example, consider the target sequence *AAB*. This sequence is non-trivial as both the within-motif dynamics and syntax is non-Markovian (Fig. 2.A). Non-Markovian sequences are generally hard to learn, because they require a memory about past dynamics (Brea et al. 2013). First, we present the target sequence repeatedly to the read-out networks (as shown in Fig. S1). After learning is finished, we test whether learning was successful by checking that the sequence is correctly produced by spontaneous dynamics (Fig. 2.B-E). Note that the slow clock spans the entire sequence (Fig. 2.C) and activates the interneurons in the correct order (Fig. 2.E), whereas the interneuron dynamics in turn determines the activation of the fast clock (Fig. 2.B) and the selection of a read-out network (Fig. 2.D). Through the learning phase, the motif weights (from the fast clock to the read-out networks) evolve to match the target motifs (Fig. 2.F), and, similarly, the syntax weights (from the slow clock to interneurons) evolve to match the ordering of the motifs in the target sequence (Fig. 2.G). Crucially, as shown below, these two sets of plastic weights are dissociated into separate pathways so that compositional sequences can be learnt efficiently through this model. Note that this conceptual model can be implemented in various ways (Appendix I) but can serve as a general framework for the learning and replay of stereotypical compositional behaviour.

**Fig. 2.**
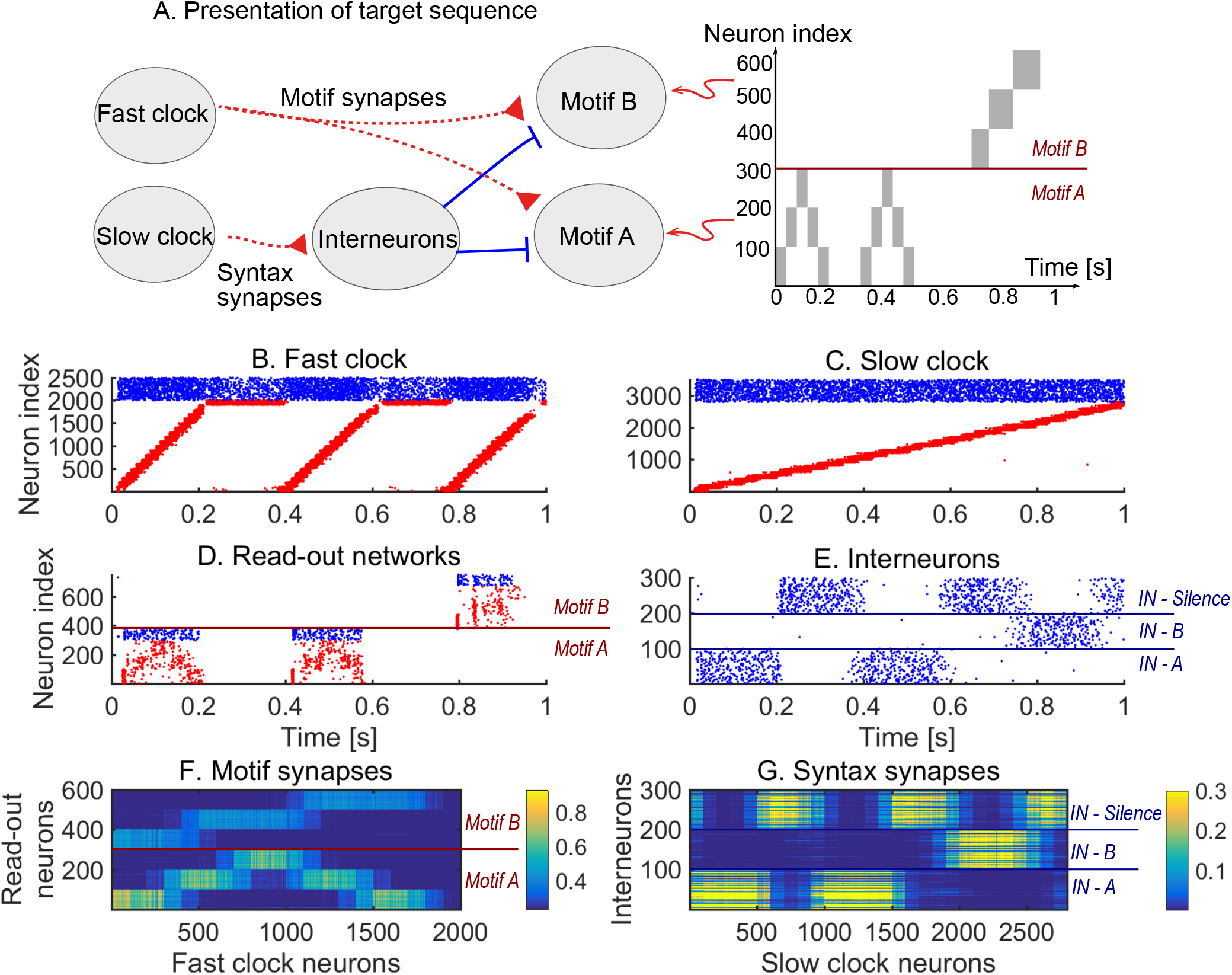
Learning sequence *AAB*. A. The target sequence is repeatedly presented to the read-out networks corresponding to motifs *A* and *B. A* and *B* are 200 ms long motifs. Between the motifs, we assume a silent period of 150 ms. B-E. Spontaneous dynamics after learning (50 target presentations). Red dots: excitatory neurons; blue dots: inhibitory neurons. B. The fast clock, controlled by interneurons 201 to 300. C. The slow clock, spanning and driving the entire sequence replay. D. The read-out networks, driven by the fast clock and controlled by the interneurons. E. The interneurons, driven by the slow clock. Neurons 1 − 100 inhibit motifs *B*. Neurons 101 − 200 inhibit motifs *A*. Neurons 201 − 300 shut down both the fast clock and read-out networks. F. The motif synapses show that the target motifs *A* (neurons 1 − 300 on the y-axis) and *B* (neurons 301 − 600 on the y-axis) are stored. The weights for motif *A* are stronger because there are two *A*s in the target sequence and only one *B*. G. The syntax weights store the temporal ordering *A*-silent-*A*-silent-*B*-silent.

### The hierarchical model enables efficient relearning of the syntax

We next demonstrate the ability of the model to relearn the ordering of the motifs. In general, we wish relearning to be efficient, i.e., the model should relearn the syntax without changing the motifs themselves. To test this idea, we perform a re-learning scheme *AAB* → *ABA* (Fig. 3). An efficient model would only learn the switch in the syntax without the need to relearn the two motifs *A* and *B*. Starting from a network where no sequence was stored, we begin with a learning phase where the sequence *AAB* is presented (as in Fig. 2) until it is learnt. We then switch to presenting the sequence *ABA* in the relearning phase. To quantify the progress of learning throughout both phases, we simulate spontaneous dynamics after every fifth target sequence presentation and compute the error between the spontaneous dynamics and the target sequence (see Methods and Fig. 3).

**Fig. 3.**
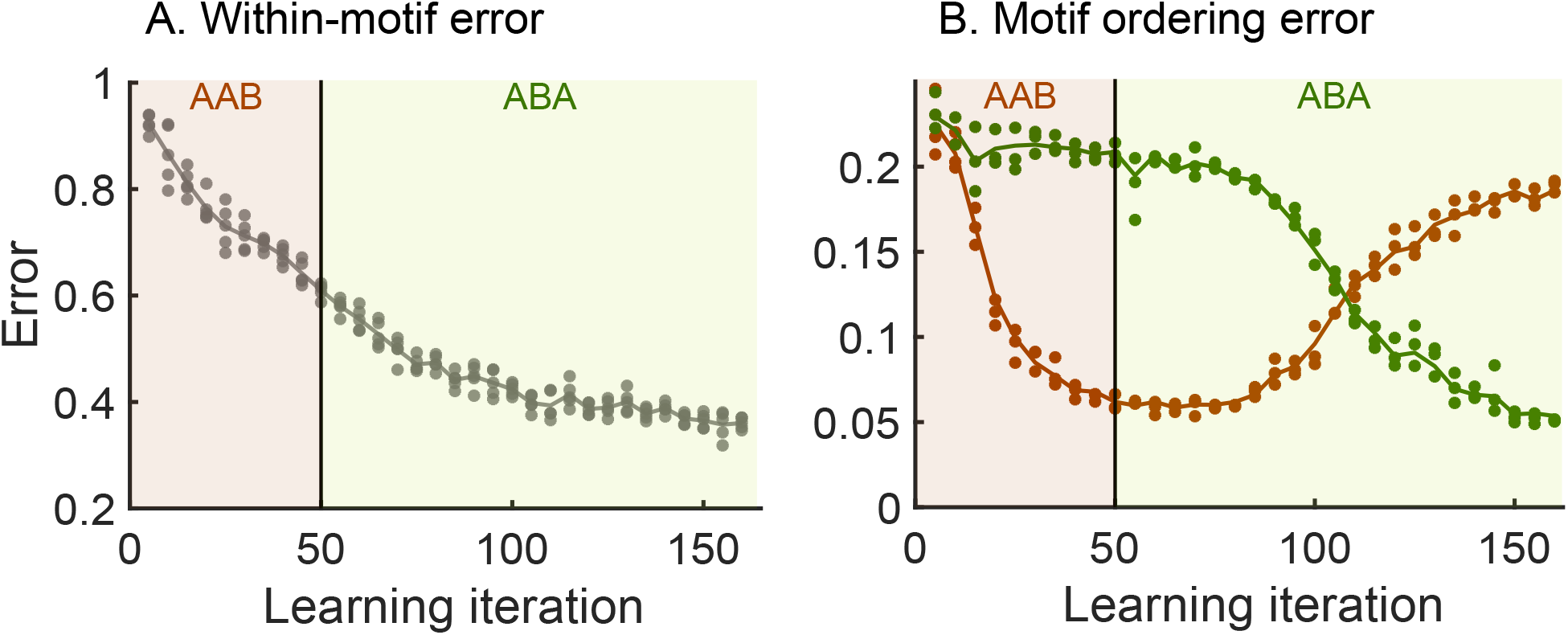
Relearning syntax: *AAB* → *ABA*. Brown shaded areas: presentation of target sequence *AAB*; dark green shaded areas: presentation of target sequence *ABA*. Brown dots: error of spontaneous dynamics with target sequence *AAB* (3 trials); dark green dots: error of spontaneous dynamics with target sequence *ABA* (3 trials). Lines guide the eye and are averages of the dots. See the methods for the details of the error measurements. A. The within-motif error keeps decreasing independent of the motif ordering. B. The motif ordering error (syntax error) switches with a delay.

Our results show that the motifs are not re-learnt when switching between the first and second target sequences—the within-motif error keeps decreasing after we switch to the relearning phase indicating that there continues to be improved learning of the motifs common to both target sequences (Fig. 3.A). In contrast, the model relearns the temporal ordering of the motifs after the switch to the new target sequence—the syntax error relative to *AAB* decreases during the learning phase and then grows during relearning at the same time as the syntax error relative to *ABA* decreases (Fig. 3.B). Therefore, the hierarchy of the model allows for efficient relearning: previously acquired motifs can be reordered into new sequences without relearning the motifs themselves.

To investigate the role of the hierarchical organisation, we next studied how the relearning behaviour compares to a *serial model* with no dissociation between motifs and syntax. The serial model contains only one clock network and the read-out networks associated with each motif, with no interneurons (Fig. S2.A). In this serial architecture, motifs and syntax are both learnt and replayed by a single pathway (Fig. S2.B and Fig. S2.C), and, consequently, when relearning the syntax, the motifs are also re-learnt from scratch even when there is no change within the individual motifs. This leads to a slower decay of the sequence error during learning and relearning in the serial model as compared to the hierarchical model (Fig. S3).

The above results illustrate the increased efficiency of the hierarchical model to learn compositional sequences. The separation of motifs and syntax into two pathways, each of them associated with a different time scale and reflected in the underlying neuronal architecture, allows for the learning and control of the different aspects of the sequence independently.

### The hierarchical organisation leads to improved learning speed and high motif fidelity

We now study the effects of having a hierarchical organisation on the speed and quality of learning. To do so, we consider three target sequences of increasing complexity, where each target sequence is comprised of a motif presented three times (Fig. 4.A).

**Fig. 4.**
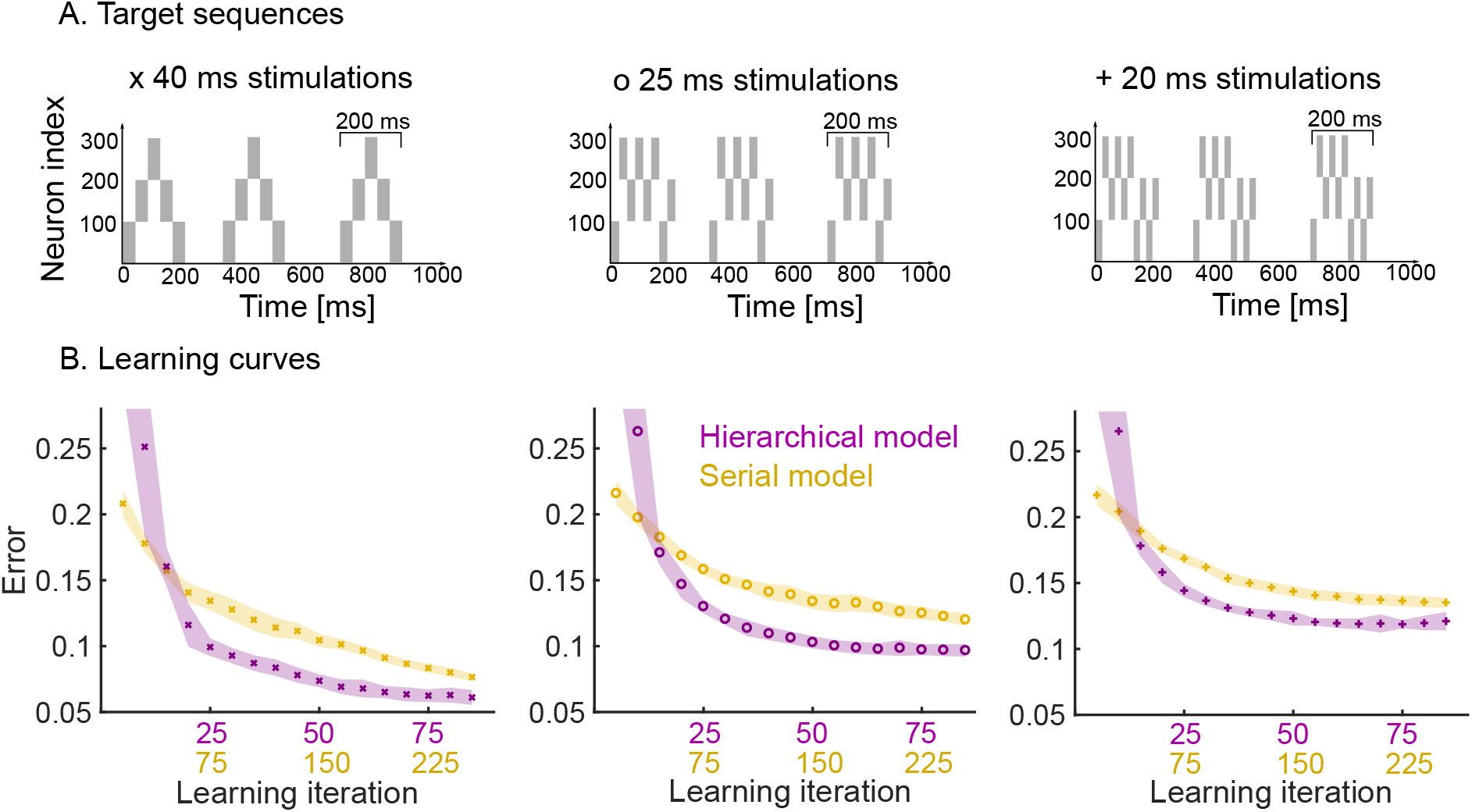
Learning speed and performance of hierarchical and serial models on three target sequences of increasing temporal complexity. A. Each target sequence consists of three presentations of the same motif (200 ms long) but with increasing complexity from left to right. Left: the simplest motif consists of five 40 ms stimulation. Middle: the motif consists of eight 25 ms stimulations. Right: the motif consists of ten 20 ms stimulations. B. Learning curves for the three target sequences for both the hierarchical and serial models. The same plasticity parameters are used for both models (see Methods). The shaded area indicates one standard deviation from the mean (50 trials). Note that the x-axis has two scales to show the three-fold increase in learning speed of the hierarchical model (i.e., for each learning iteration of the hierarchical model there are three iterations of the serial model). The performance degrades from left to right, as a more difficult target sequence is presented.

First, we studied the speed at which the pattern is learnt by the hierarchical model as compared to the serial model. The hierarchical model is roughly three times faster than the serial model in learning patterns consisting of three repetitions (Fig. 4.B). This is expected: in the hierarchical model, the same motif synapses are potentiated three times during a single target presentation, whereas no such repeated learning takes place in the serial model. Furthermore, the speed and quality of the learning also depends on the complexity of the target sequence, i.e., target sequences with rapid temporal changes are harder to learn. Learning target sequences with faster-changing, more complex temporal features leads to a degradation of the performance of both models, but the hierarchical model consistently learns roughly three times faster than the serial model for all patterns (Fig. 4, left to right).

Another important quality measure of learning is the reliability and consistency of the pattern replayed by the model under spontaneous dynamics. To study this, we generated repeated trials in which the three target sequences learnt (in Fig. 4) were replayed spontaneously, and we compared the variability of the read-out dynamics across the trials for both the hierarchical and serial models. We first computed the within-motif and between-motif variability in the spontaneous trials. The hierarchical model leads to low within-motif variability and higher variability in the between-motif timings. This follows from the spiking in the read-out networks, with highly variable time gaps between motifs in the hierarchical model (Fig. 5.A). On the other hand, the spike trains within the three motifs correlate strongly with each other for the hierarchical model (Fig. 5.B). This is the case for the three target sequences.

**Fig. 5.**
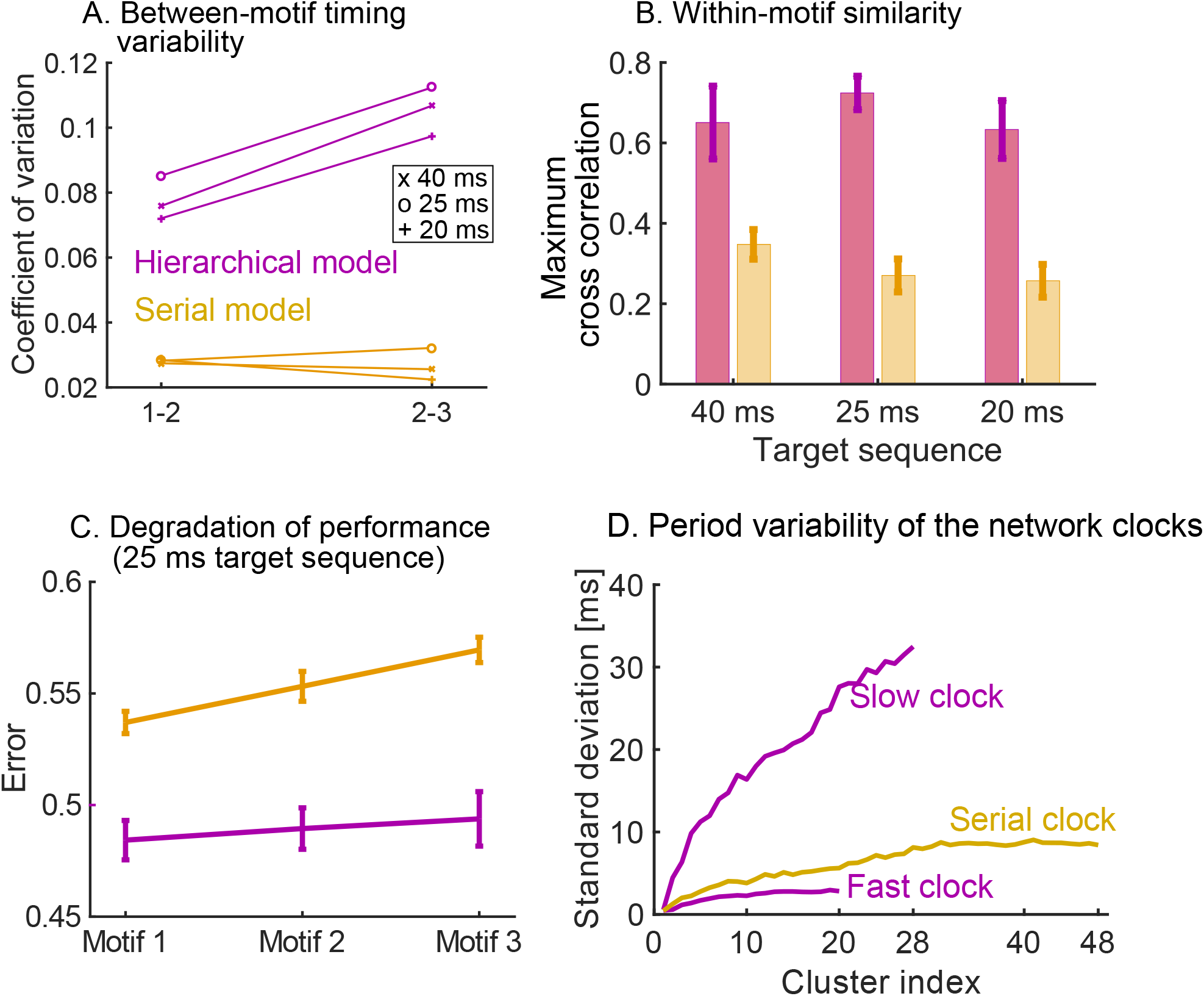
Measuring variability and performance in the read-out dynamics. A. The time between motifs 1 and 2 and motifs 2 and 3 is measured during spontaneous dynamics. We plot the coefficient of variation of these times (50 trials) on the y-axis, for the three target sequences in Fig. 4.A. B. The cross correlation between the spike trains in the first motif and the second and third motif is measured, normalized by the auto-correlation of the spike trains in motif 1. The maximum of the cross correlation is recorded in each trial (50 trials). This is repeated for the three target sequences in Fig. 4.A. C. We measure the error between the target sequence with 25 ms stimulations in Fig. 4.A and spike trains in motif 1, 2 and 3. In both models, the performance degrades towards later occurring motifs. The degradation is significantly worse in the serial model: a linear regression yields a slope of 0.0163 for the serial model and a slope of 0.0048 for the hierarchical model (*p <* 10^−5^ using t-test). D. The serial clock (48 clusters) is obtained by adding the slow (28 clusters) and fast (20 clusters) clocks together. Sequential dynamics is simulated 50 times for each clock. The time at which each cluster is activated in the sequential dynamics is measured. The standard deviation of these activation times is plotted as a function of the cluster index. The serial clock has a maximal variability of about 9 ms. The fast and slow clock have a maximal variability of about 3 and 35 ms respectively.

We then studied the consistency of the motif as it is repeated (three times) within a target sequence. We observe that a high degradation of the repeated motif towards the end of the sequence in the serial model, which is milder in the hierarchical model (Fig. 5.C). In summary, the hierarchical model produces accurate motifs that persist strongly over time, but with higher variability in the timing between them. The high reliability of the motifs is due to the stronger learning on the motif synapses discussed above. The higher variability in the inter-motif times is a result of the underlying variability of the periods of the clock networks. As discussed in Ref. Maes et al. 2020, the sequential dynamics that underpins the clock networks operates by creating clusters of neurons that are active over successive periods of time. In that sense, the network uses neuron clusters to discretise a span of time (its period) into time increments. The variability of the period of the clock depends on the number of clusters, the number of neurons per cluster in the network, and the time span to discretise. A fast clock will thus have low variability in its period, whereas the slow clock is highly variable. The variability of the period of the serial clock is between the fast and slow clocks (Fig. 5.D). Consequently, within-motif temporal accuracy is maintained quasi-uniformly over the sequence in a hierarchical model. The price to pay is the addition of appropriately wired interneurons.

### The hierarchical organisation increases the capacity of the model to store different sequences

As shown above, the plasticity of the model allows it to relearn single sequences, yet the relearning process might be too slow for particular situations. In general, animals acquire and store multiple sequences to be used as needed. Motivated by this idea, we explore the capacity of the hierarchical model to learn, store and replay more than one sequence, and we compare it to the alternative serial model. First, we note that a new sequence can be stored in the hierarchical model by adding another interneuron network in parallel. The additional interneuron network is a replica of the existing one, with the same structure and connection weights to the rest of the system.

Each interneuron network learns one syntax, in the same way as one read-out network learns one motif. As an illustration, we learn the sequences *AAB* and *BAAB* (Fig. 6.A), by presenting the target sequences alternately. We then simulate spontaneous dynamics to test that the learning is successful. The spiking dynamics (Fig. 6.B-E) show that the model is able to replay the two sequences. To select between the two sequences, we use an attentional external current to the interneuron networks during learning and spontaneous dynamics (shaded areas in Fig. 6.E). Depending on the interneuron activity, the fast clock (Fig. 6.B) and read-out networks (Fig. 6.D) are active. Note that the motifs are encoded in the motif weights (Fig. 6.F) and syntax weights encode both target motif orderings (Fig. 6.G). These results show that the hierarchical model can learn, store and replay multiple sequences. Importantly, the motifs are still flexibly re-used: when motifs *A* and *B* are learnt by presenting sequence *AAB*, they can immediately be re-used when a different syntax (e.g., *BAAB*) is presented.

**Fig. 6.**
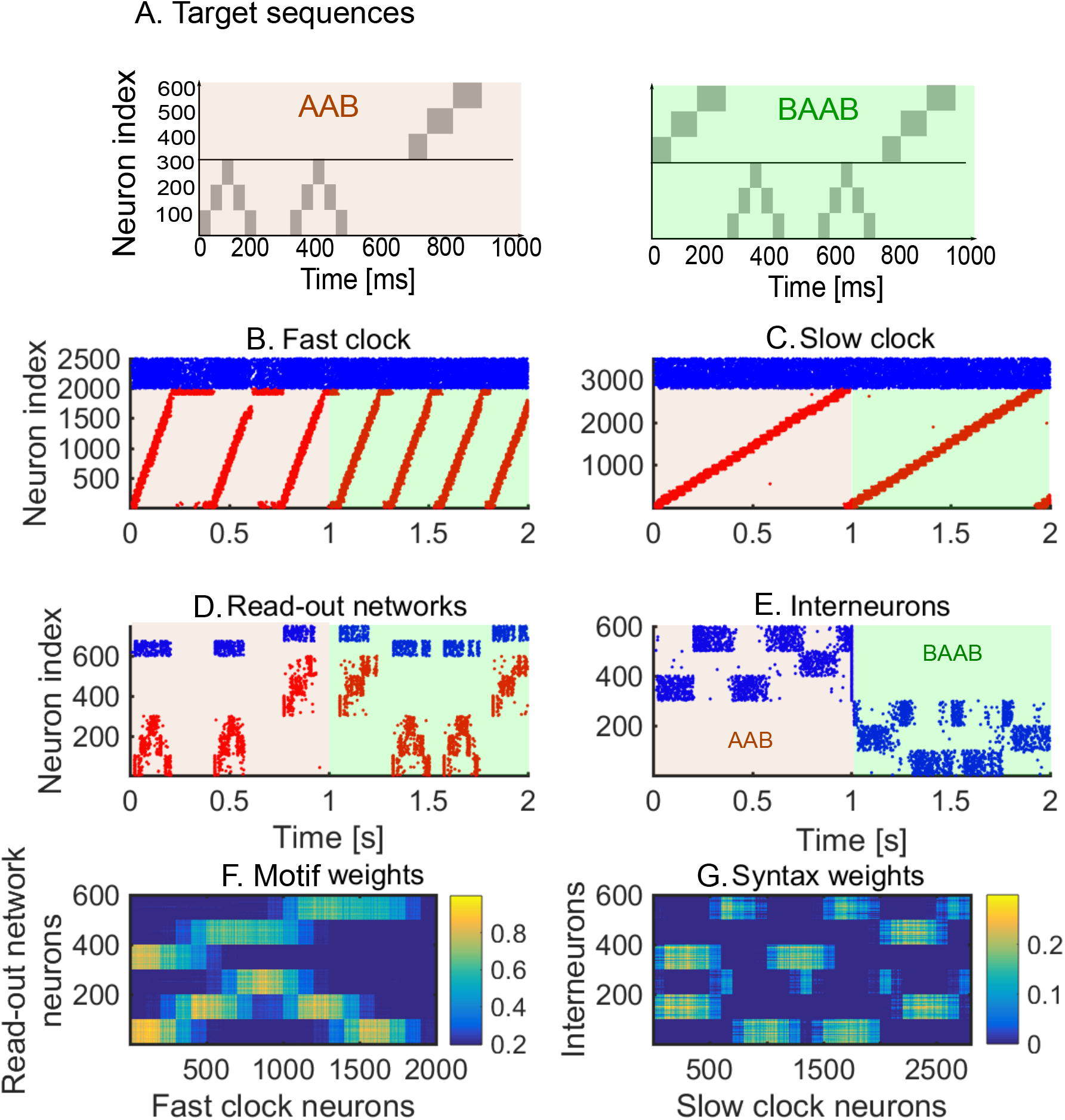
Spontaneous dynamics after learning two sequences alternately (80 learning iterations). A. The target sequences. B-E. Red dots: excitatory neurons; blue dots: inhibitory neurons. Brown shaded area: sequence *AAB* is played by inhibiting the interneurons related to the second sequence; light green shaded area: sequence *BAAB* is played by inhibiting the interneurons related to the first sequence. B. Spike raster of the fast clock. C. Spike raster of the slow clock. D. Spike raster of the two read-out networks. E. Spike raster of the interneurons. An external attentional inhibitory current selects which sequence is played. F. The motif weights encode the two motifs. Note the similarity with Fig. 2.F: the same motifs are re-used in both sequences. G. The syntax weights encode the two motif orderings. Note the difference with Fig. 2.G: an additional syntax is stored. All motif and syntax synapses are plastic at all times during the sequence presentations.

We then compare the capacity of the hierarchical model to the serial model (Fig. S4). In the serial model, read-out networks have to be added in order to learn and store multiple sequences. This is inefficient for two reasons: 1) The same motif might be stored in different read-out networks; The addition of new read-out networks in the serial model requires more ‘hardware’ (i.e. neurons and synapses) than the addition of an interneuron network in the hierarchical model. Our results show that the hierarchical model can learn and store multiple sequences by adding more interneuron networks in parallel. This hierarchical organisation thus exploit the compositional nature of the sequences in a way the serial model cannot leading to increased capacity.

### The hierarchical model displays increased robustness to perturbations of the sequential dynamics

We next investigate the role of the hierarchy in enhancing the robustness of the model to perturbations in the target sequence. Behavioural perturbation experiments have shown that individual motifs can be removed mid-sequence without affecting later-occurring motifs (Geddes et al. 2018). This is a useful feature which can dramatically improve the robustness of responses, since later-occurring behaviour does not critically depend on the successful completion of all previous behaviours. To examine this issue, we have carried out simulations on the serial and hierarchical models under perturbations in the firing of the neurons in the clock; specifically we remove the external input to excitatory neurons in the clock network. In the serial model, the perturbation leads to the breakdown of the remaining parts of the sequence (Fig. 7.A), whereas when the same perturbation is applied to the fast clock of the hierarchical model, we see that later motifs are preserved (Fig. 7.B). The reason is that the dynamics in the slow clock is intact and continues to drive the behaviour. Perturbing the slow clock, and keeping the fast clock intact, has less predictable outcomes for the dynamics. Random activity in the interneurons can cause motifs to be played in a random order (Fig. S5). Overall, the hierarchical model improves the robustness. Indeed, at any point in time, a single cluster of neurons is active in the clock of the serial model, whereas there are two active clusters of neurons (one in the fast clock and another in the slow clock) in the hierarchical model. This separation of time scales is fundamental to preserve the robustness of the model.

**Fig. 7.**
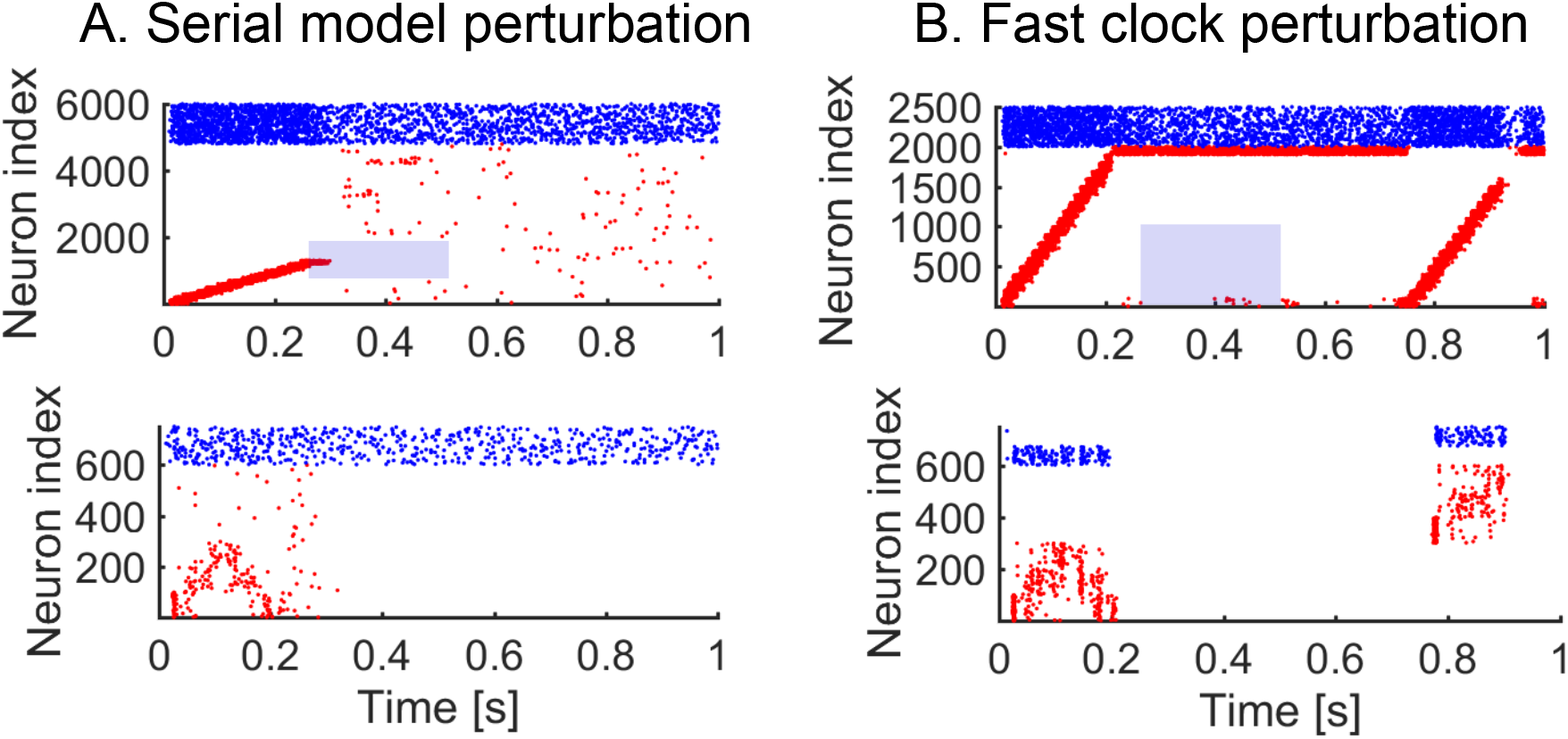
Perturbing the dynamics. We learn sequence *AAB* and then apply a perturbation. Blue shade indicates the perturbation time, and neurons perturbed. A. 250 ms perturbation of the serial network clock. The targeted neurons (neurons 1000 to 2000) have no excitatory external input during the perturbation. The sequential activity breaks down completely. B. 250 ms perturbation of the fast clock in the hierarchical model. The targeted neurons (neurons 1 to 1000) have no excitatory external input during perturbation. The sequential activity breaks down but is reactivated for the final motif through the interneurons.

## DISCUSSION

### Summary of results

We have presented here a hierarchical neuronal network model for the learning of compositional sequences. We demonstrated how motifs and syntax can be learnt independently of each other. The hierarchical structure has direct implications for the learning and is contrasted with a serial architecture. The hierarchical structure leads to an increased learning speed and the possibility to efficiently relearn the ordering of individual motifs. The replays of individual motifs are more similar to each other as compared to replays in the serial model. Separating the motifs and syntax into two different pathways in the hierarchical model has also implications for the capacity and robustness. The motifs can be re-used in the hierarchical model, leading to a significantly higher capacity when the number of interneurons is much smaller than the number of read-out neurons. Finally, the serial model has a single pathway, as opposed to two, and is therefore more prone to perturbations.

### From serial to hierarchical modelling

Modelling studies so far have either focused on the study of sequential dynamics (Chenkov et al. 2017; Billeh and Schaub 2018; Setareh et al. 2018; Spreizer et al. 2019) or on motif acquisition (Stroud et al. 2018; Logiaco et al. 2019; Maes et al. 2020). This paper introduces an explicitly hierarchical model as a fundamental building block for the learning and replay of sequential dynamics of a compositional nature. Sequential dynamics is omnipresent in the brain and might be important in time-keeping during behaviour (Hahnloser et al. 2002; Ikegaya et al. 2004; Harvey et al. 2012; Peters et al. 2014; Katlowitz et al. 2018; Adler et al. 2019). When temporal sequences are compositional (i.e., formed by the ordering of motifs), they lead to the presence of different time scales associated with the motifs and their ordering (or syntax). From the perspective of learning, such multiscale temporal organisation lends itself naturally to a hierarchical organisation, where the different scales are associated with different groups of neurons in the network (see also Schaub et al. 2015).

Hierarchical temporal structures might arise during development in a variety of ways (Dominici et al. 2011; Yang et al. 2019). One way is that a single *proto*sequence is learnt first. The *proto*sequence covers the entire behaviour learning the most crude aspects. This might then be followed by splitting the *proto*sequence into multiple sequences specialized to different aspects of the behaviour. A similar splitting of sequences has been observed in birdsong (Fiete et al. 2010; Okubo et al. 2015). Hierarchical motor control has also been studied in the artificial intelligence field (Merel et al. 2019). A recent model works towards closing the gap from a machine system to a biological system (Logiaco and Escola 2020) but remains non-trivial to implement using dynamics and plasticity that are considered to be more realistic in a biological sense.

### A storage and replay device

The proposed model can be viewed as a biological memory device that stores sequences by means of supervised learning and replays them later by activating the device with spontaneous activity. However, it is important to note that during spontaneous activity there is no input to the device other than the random spike patterns that keep the dynamics of the system going. This mode of operation is therefore distinct from computational machines, such as the liquid state machine (Maass et al. 2002; Maass 2011) or the tempotron (Gütig and Sompolinsky 2006), where different input patterns are associated with and transformed into different output patterns. Such computational machines, where spiking patterns are inputs to be transformed or classified, are thus complementary to our autonomous memory device.

### Hierarchies in other tasks

Hierarchies exist beyond the somewhat simple learning of compositional sequences, and it is expected that hierarchical models share common basic features despite solving highly distinct problems. For instance, a recent example of a hierarchical model for working memory uses two different networks: an associative network and a *task-set* network (Bouchacourt et al. 2020). In our setting, the associative network could be identified with the motifs (fast clock+read-out) whereas the *task-set* network would correspond to the syntax (slow clock+interneurons). Navigation is another typical example of a task where hierarchy is used (Tomov et al. 2020), and the discovery of structure in an environment is closely related to the presence of a hierarchy (Karuza et al. 2016).

### Relating the model to experiments

As mentioned above, there are *qualitative* similarities between the proposed hierarchical model and experimental studies. Experimental studies have pointed increasingly at the importance of hierarchical organisation both in structural studies as well as in the learning and execution of movement and auditory sequences. For example, behavioural re-learning has shown that birds can re-order motifs independently from the within-motif dynamics (Lipkind et al. 2017). Optogenetical perturbation in the striatum of mice has shown that individual motifs can be deleted or inserted mid-sequence, without altering the later part of the behavioural sequence (Geddes et al. 2018). The proposed model aims to provide a conceptual framework to explain such behavioural observations while simultaneously using biophysically realistic spiking networks and plasticity rules.

However, a *quantitative* link between model and experiment is not trivial. This is true for behaviour, but even more so for neural activity. Indeed, our model has free parameters, including topology and plasticity, which need to be tuned to the task at hand. Nevertheless, there are two recent advances that may help future work in this direction. Firstly, there have been recent technological improvements in recording of behaviour (Egnor and Branson 2016; Berman 2018) and neural activity (Jun et al. 2017) along with the possibility to apply perturbations (Geddes et al. 2018). Secondly, there has been progress in decoding meaningful information from large observational datasets (Williams et al. 2018), e,g„ the extraction of sequences from neural recordings (Mackevicius et al. 2019) and the analysis of learning behaviour of songbirds (Kollmorgen et al. 2020). In this vein, an interesting question to pursue is whether one could *rediscover* the hierarchical structure from temporal data generated by our model. For instance, one could observe a randomly chosen subset of neurons in the model: could the hierarchical organisation and function of the network be inferred from those partial observations by using data analysis?

## Conclusion

Using realistic plasticity rules, we built a spiking network model for the learning of compositional temporal sequences of motifs over multiple time scales. We showed that a hierarchical model is more flexible, efficient and robust than the corresponding serial model for the learning of such sequences. The hierarchical model concentrates the variability in the inter-motif timings but achieves high motif fidelity.

## METHODS AND MATERIALS

Excitatory neurons (*E*) are modelled with the adaptive exponential integrate-and-fire model (Brette and Gerstner 2005). A classical integrate-and-fire model is used for the inhibitory neurons (*I*). Motifs and syntax are learnt using simple STDP-rules (see for example Kempter et al. 1999) without need for additional fast normalization mechanisms.

### Model architecture

The hierarchical model consists of four recurrent networks. Each network and their parameters are described below. Synaptic weights within each recurrent network are non-zero with probability *p* = 0.2. The synaptic weights in the recurrent networks which produce sequential dynamics are scaled using a scaling factor 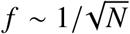, i.e., it scales with the corresponding network size *N*.

### Fast clock (Fc)

The fast clock network has 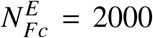 excitatory and 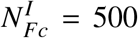 inhibitory neurons recurrently connected with parameters shown in Table 1. Sequential dynamics is ensured by dividing the excitatory neurons in 20 clusters of 100 neurons. The baseline excitatory weights 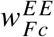 within the same cluster are multiplied with a factor of 25, whereas the excitatory weights from cluster *i* to cluster *i* + 1 mod 20 (*i* = 1..20) are multiplied by a factor of 12.5. Previous studies have shown that such a weight structure leads to sequential dynamics and can be learnt in a biologically plausible way (Zheng and Triesch 2014; Murray and Escola 2017; Maes et al. 2020). The fast clock receives excitatory external random Poisson input and inhibitory input from the interneurons, and projects to the read-out networks.

**TABLE 1.** Fast clock network parameters

| Constant | Value | Description |
| --- | --- | --- |
| $N_{Fc}^E$ | 2000 | Number of recurrent E neurons |
| $N_{Fc}^I$ | 500 | Number of recurrent I neurons |
| $f$ | 0.6325 | Scaling factor |
| $w_{Fc}^{EE}$ | $5f$ pF | Baseline E to E synaptic strength |
| $w_{Fc}^{IE}$ | $3.5f$ pF | E to I synaptic strength |
| $w_{Fc}^{EI}$ | $110f$ pF | I to E synaptic strength |
| $w_{Fc}^{II}$ | $36f$ pF | I to I synaptic strength |

### Read-out networks (R)

Each read-out network codes for one individual motif. There are no overlaps or connections between the different read-out networks. The read-out networks are identical and balanced (see Table 2 for the parameters). The read-out networks receive excitatory input from the fast clock and inhibitory input from the interneurons. They also receive external inputs: a supervisor component (only during learning) and a random input.

**TABLE 2.**
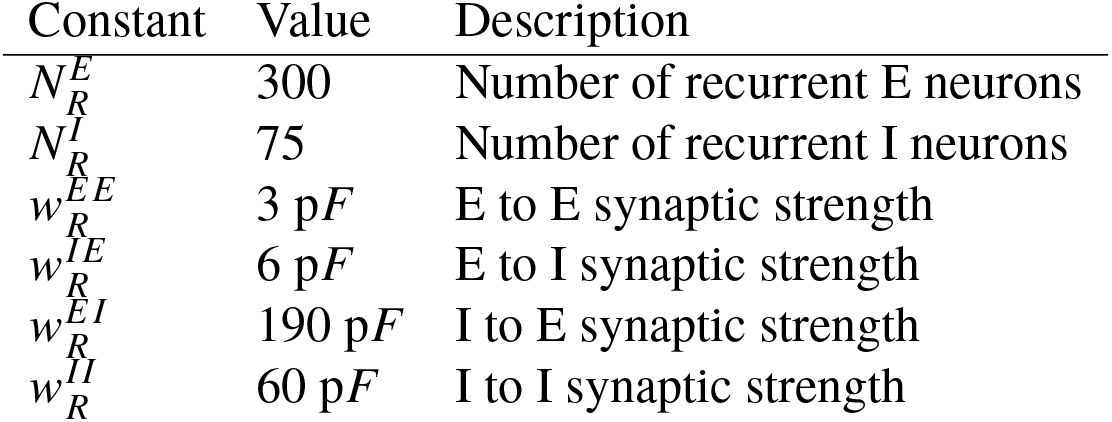
Read-out network parameters

### Slow clock (Sc)

The slow clock network has 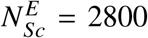 excitatory and 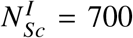 inhibitory neurons, recurrently connected. It is essentially a scaled copy of the fast clock. Table 3 shows the parameters of this network. Sequential dynamics is ensured by dividing the excitatory neurons in 28 clusters of 100 neurons. The baseline excitatory weights 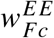 within the same cluster are multiplied with a factor of 25, and the excitatory weights from cluster *i* to cluster *i* + 1 mod 28 (*i* = 1..28) are multiplied by a factor of 4.7. The slow clock receives excitatory external random Poisson input and projects to the interneuron networks.

**TABLE 3.** Slow clock network parameters

| Constant | Value | Description |
| --- | --- | --- |
| $N_{Sc}^E$ | 2800 | Number of recurrent E neurons |
| $N_{Sc}^I$ | 700 | Number of recurrent I neurons |
| $f$ | 0.5345 | Scaling factor |
| $w_{Sc}^{EE}$ | 5f pF | Baseline E to E synaptic strength |
| $w_{Sc}^{IE}$ | 3.5f pF | E to I synaptic strength |
| $w_{Sc}^{EI}$ | 110f pF | I to E synaptic strength |
| $w_{Sc}^{II}$ | 36f pF | I to I synaptic strength |

### Interneuron networks (In)

Each interneuron network codes for one syntax. There are no overlaps between the interneuron networks. Each interneuron network is balanced with parameters given in Table 4. Neurons within each interneuron network are grouped into 3 groups of 100 neurons: one group per motif and one group for the ‘silent’ motif. The interneuron networks receive excitatory input from all other networks. They also receive random excitatory external input.

**TABLE 4.** Interneuron network parameters

| Constant | Value | Description |
| --- | --- | --- |
| $N_{In}^I$ | 300 | Number of recurrent I neurons |
| $w_{In}^{II}$ | 25 pF | I to I synaptic strength |

### Connections between recurrent networks

The recurrent networks are connected to each other to form the complete hierarchical architecture. All excitatory neurons from the fast clock project to all excitatory neurons in the read-out networks. These synapses, 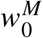, are plastic. All excitatory neurons from the slow clock project to all the interneurons. These synapses, 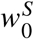, are also plastic. To signal the end of a motif, the penultimate cluster of the fast clock activates the interneurons of the ‘silent’ motif. The last cluster is also connected to the ‘silent’ motif which silences all other clusters in the fast clock and all neurons in the read-out networks. Each read-out network gives excitatory input to its corresponding interneuron group. This interneuron group laterally inhibits the other read-out network(s). Table 5 gives all the parameters of the connections between the different networks.

**TABLE 5.**
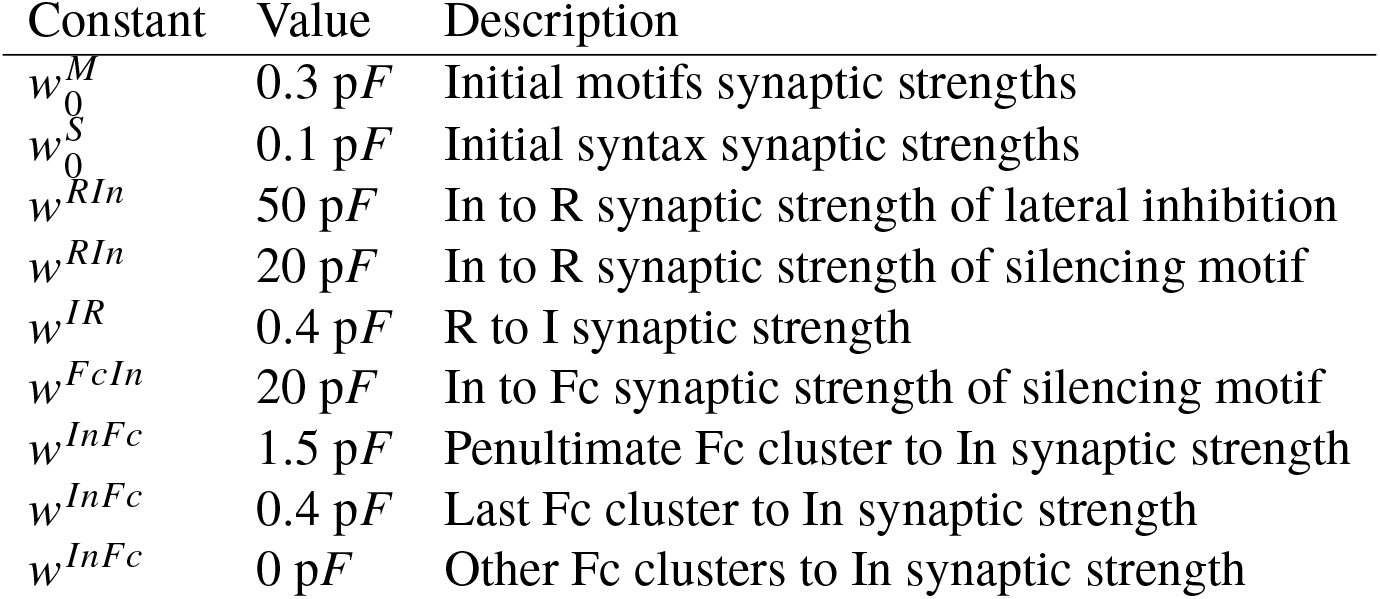
Connections between four networks

### Serial model (Sm)

The hierarchical model is compared with a serial model. The serial model has one large clock (with the same number of neurons as the fast and slow clocks combined) and no interneurons. Sequential dynamics is generated by clustering the neurons in the network in 48 clusters of 100 neurons. The baseline excitatory weights 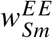 of the same cluster are multiplied with a factor of 25, and the excitatory weights from group *i* to group *i* + 1 mod 48 (*i* = 1..48) are multiplied by a factor of 6. Table 6 shows the network parameters. The read-out network is kept unchanged (Table 2).

**TABLE 6.**
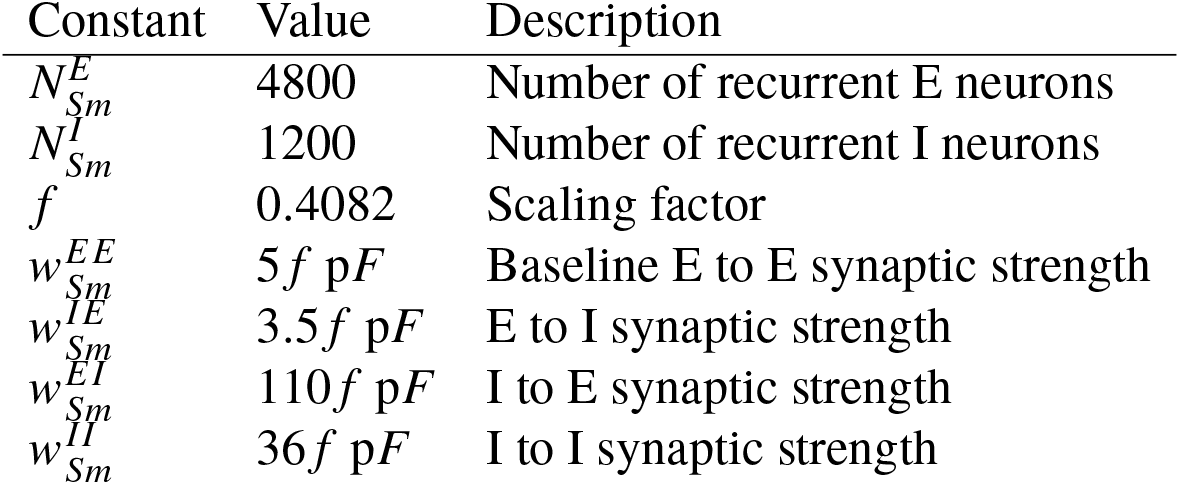
Serial model clock network parameters

### Neural and synaptic dynamics

All neurons in the model are either excitatory (*E*) or inhibitory (*I*). The parameters of the neurons do not change depending on which network they belong to. Parameters are consistent with Ref. Litwin-Kumar and Doiron 2014.

### Membrane potential dynamics

The membrane potential of the excitatory neurons (*V*^*E*^) has the following dynamics:

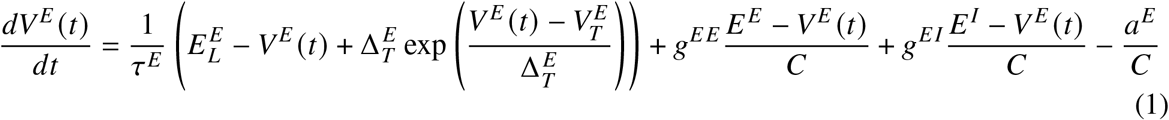

where τ^*E*^ is the membrane time constant, 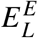 is the reversal potential, 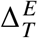 is the slope of the exponential, *C* is the capacitance, *g*^*EE*^, *g*^*EI*^ are synaptic input from excitatory and inhibitory neurons respectively and *E*^*E*^, *E*^*I*^ are the excitatory and inhibitory reversal potentials respectively. When the membrane potential diverges and exceeds 20 mV, the neuron fires a spike and the membrane potential is reset to *V*_*r*_. This reset potential is the same for all neurons in the model. There is an absolute refractory period of τ_*ahs*_. The parameter 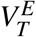 is adaptive for excitatory neurons and set to *V*_*T*_ + *A*_*T*_ after a spike, relaxing back to *V*_*T*_ with time constant τ_*T*_ :

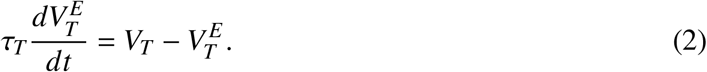

The adaptation current *a*^*E*^ for excitatory neurons follows:

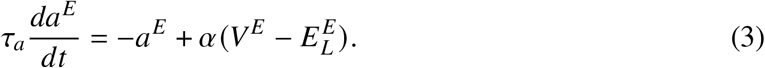

where τ_*a*_ is the time constant for the adaptation current. The adaptation current is increased with a constant *β* when the neuron spikes.

The membrane potential of the inhibitory neurons (*V* ^*I*^) has the following dynamics:

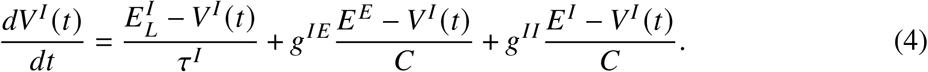

where τ^*I*^ is the inhibitory membrane time constant, 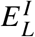 is the inhibitory reversal potential and *E*^*E*^, *E*^*I*^ are the excitatory and inhibitory resting potentials respectively. *g*^*EE*^ and *g*^*EI*^ are synaptic input from excitatory and inhibitory neurons respectively. Inhibitory neurons spike when the membrane potential crosses the threshold *V*_*T*_, which is non-adaptive. After this, there is an absolute refractory period of τ_*ahs*_. There is no adaptation current (see Table 7 for the parameters of the membrane dynamics).

**TABLE 7.** Neuronal membrane dynamics parameters

| Constant | Value | Description |
| --- | --- | --- |
| $\tau_E$ | 20 ms | E membrane potential time constant |
| $\tau_I$ | 20 ms | I membrane potential time constant |
| $\tau_{abs}$ | 5 ms | Refractory period of E and I neurons |
| $E^E$ | 0 mV | excitatory reversal potential |
| $E^I$ | -75 mV | inhibitory reversal potential |
| $E_L^E$ | -70 mV | excitatory resting potential |
| $E_L^I$ | -62 mV | inhibitory resting potential |
| $V_r$ | -60 mV | Reset potential (both E and I) |
| $C$ | 300 pF | Capacitance |
| $\Delta_T^E$ | 2 mV | Exponential slope |
| $\tau_T$ | 30 ms | Adaptive threshold time constant |
| $V_T$ | -52 mV | Membrane potential threshold |
| $A_T$ | 10 mV | Adaptive threshold increase constant |
| $\tau_a$ | 100 ms | Adaptation current time constant |
| $\alpha$ | 4 nS | Adaptation current factor |
| $\beta$ | 0.805 pA | Adaptation current increase constant |

### Synaptic dynamics

The synaptic conductance, *g*, of a neuron *i* is time dependent, it is a convolution of a kernel with the total input to the neuron *i*:

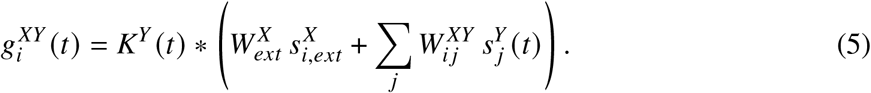

where *X* and *Y* can be either *E* or *I. K* is the difference of exponentials kernel:

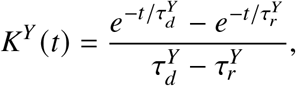

with a decay time τ_*d*_ and a rise time τ_*r*_ dependent only on whether the neuron is excitatory or inhibitory. The conductance is a sum of recurrent input and external input. The externally incoming spike trains 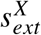 are generated from a Poisson process with rates 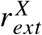. The externally generated spike trains enter the network through synapses 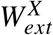 (see Table 8 for the parameters of the synaptic dynamics).

**TABLE 8.** Synaptic dynamics parameters

| Constant | Value | Description |
| --- | --- | --- |
| $\tau_d^E$ | 6 ms | E decay time constant |
| $\tau_r^E$ | 1 ms | E rise time constant |
| $\tau_d^I$ | 2 ms | I rise time constant |
| $\tau_r^I$ | 0.5 ms | I rise time constant |
| $W_{ext}^E$ | 1.6 pF | External input synaptic strength to E neurons |
| $r_{ext}^E$ | 4.5 kHz | Rate of external input to E neurons |
| $W_{ext}^I$ | 1.52 pF | External input synaptic strength to I neurons |
| $r_{ext}^I$ | 2.25 kHz | Rate of external input to I neurons |

### Plasticity

#### Motif plasticity

The synaptic weight from excitatory neuron *j* in the fast clock network to excitatory neuron *i* in the read-out network is changed according to the following differential equation:

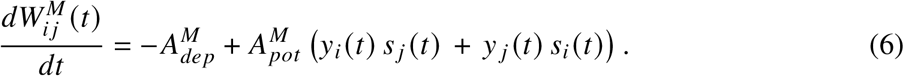

where 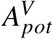 and 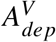 are the amplitude of potentiation and depression, *s*_*i*_ (*t*) is the spike train of the postsynaptic neurons, and *s*_*j*_ (*t*) is the spike train of the presynaptic neurons. Both pre- and post-synaptic spike trains are low pass filtered with time constant τ_*M*_ to obtain *y* (*t*):

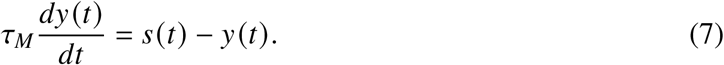

The synapses from the fast clock to the read-out network have a lower and upper bound 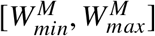. Table 9 shows parameter values for the motif plasticity rule.

**TABLE 9.** Motif plasticity parameters

| Constant | Value | Description |
| --- | --- | --- |
| $A_{Mpot}$ | 0.03 pFHz | Amplitude of potentiation |
| $A_{Mdep}$ | $2/3 \times 10^{-6}$ pF | Amplitude of depression |
| $\tau_M$ | 5 ms | Time constant of low pass filter |
| $W_{min}^M$ | 0 pF | Minimum I to I weight |
| $W_{max}^M$ | 1 pF | Maximum I to I weight |

#### Syntax plasticity

Similar to the motif plasticity rule, the syntax plasticity rule has a symmetric window. The dynamics is as such governed by the same equations, with slightly different parameters:

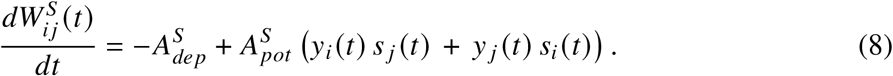

where *s*_*i*_ (*t*) is the spike train of the postsynaptic neurons, and *s* _*j*_ (*t*) is the spike train of the presynaptic neurons. The spike trains are low pass filtered with time constant τ_*S*_ to obtain *y* (*t*) (as in equation 7). The synapses from the slow clock to the interneurons have a lower and upper bound 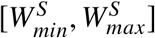. Table 10 shows parameter values for the syntax plasticity rule. Note that the time constants are longer than the time constants in the motif plasticity.

**TABLE 10.**
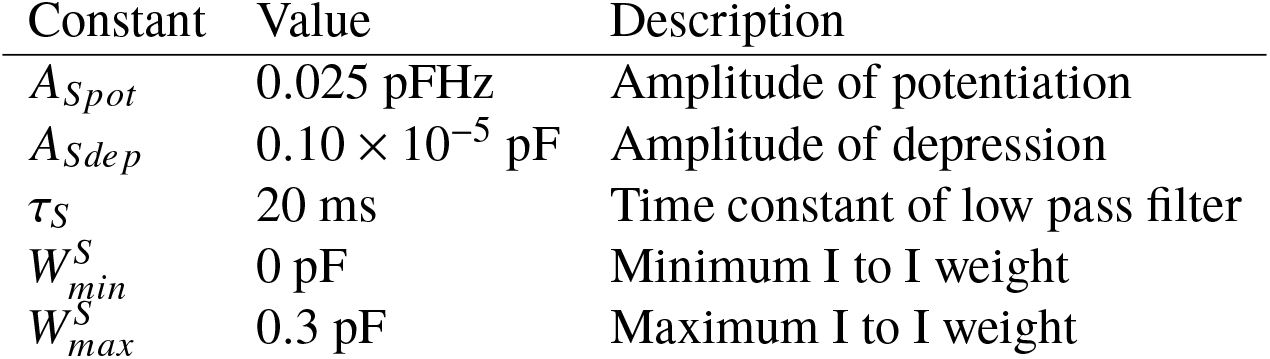
Syntax plasticity parameters

### Measuring the error

#### Motif and total error

The spontaneous dynamics of the excitatory read-out neurons is compared to a binary target sequence to measure the error during learning. The target sequence is the sequence of motifs or a single motif depending on which error is computed. The spontaneous spiking dynamics is first convolved using a Gaussian kernel of width ∼ 10 ms. This gives a proxy to the firing rates of the neurons. The firing rates are then normalized between 0 and 1. Dynamic time warping is finally used to compare the normalized spontaneous dynamics to the target sequence. Dynamic time warping is needed to remove the timing variability in the spontaneous dynamics. We computed dynamic time warping using the built-in Matlab function *dtw*. Dynamic time warping was not used to compute the error in Fig. 5.

#### The ordering error

The target sequence is now the binary target dynamics of the interneurons. Similarly as before, the spontaneous dynamics of the interneurons is convolved and normalized to compute the error with the target using dynamic time warping.

### Numerical simulations

#### Protocol – learning

A start current of 5 kHz is given for 10 ms to the first cluster of the slow clock to initiate a training session. Strong supervising input (50 kHz) to the read-out networks controls the dynamics in the read-out networks. The weights from the read-out networks to the interneurons make sure that also the interneurons follow the target ordering: there is no need for an explicit target current to the interneurons. At the start of each motif the fast clock is activated by giving a strong current of 50 kHz to the first cluster for 40 ms. The high supervisor currents are assumed to originate from a large network of neurons, external to this model.

#### Protocol – spontaneous dynamics

A start current of 5 kHz is given for 10 ms to the first cluster of the slow clock to initiate a spontaneous replay. The slow clock determines which interneurons are active, together with an external attention mechanism (if multiple sequences are stored). The interneurons then determine which read-out network is active. The fast dynamics in the read-out networks is controlled by the input from the fast clock.

#### Simulations

The code used for the training and testing of the spiking network model is built in Matlab. Forward Euler discretisation with a time step of 0.1 ms is used. The code will be made available after publication.

## ACKNOWLEDGMENTS

We thank Victor Pedrosa and Barbara Feulner for helpful comments on the manuscript. AM acknowledges funding through the EPSRC Centre for Neurotechnology. MB acknowledges funding through EPSRC award EP/N014529/1 supporting the EPSRC Centre for Mathematics of Precision Healthcare at Imperial. CC acknowledges support by BBSRC BB/N013956/1, BB/N019008/1, Wellcome Trust 200790/Z/16/Z, Simons Foundation 564408 and EPSRC EP/R035806/1.

## SUPPLEMENTARY FIGURES

**Fig. S1.**
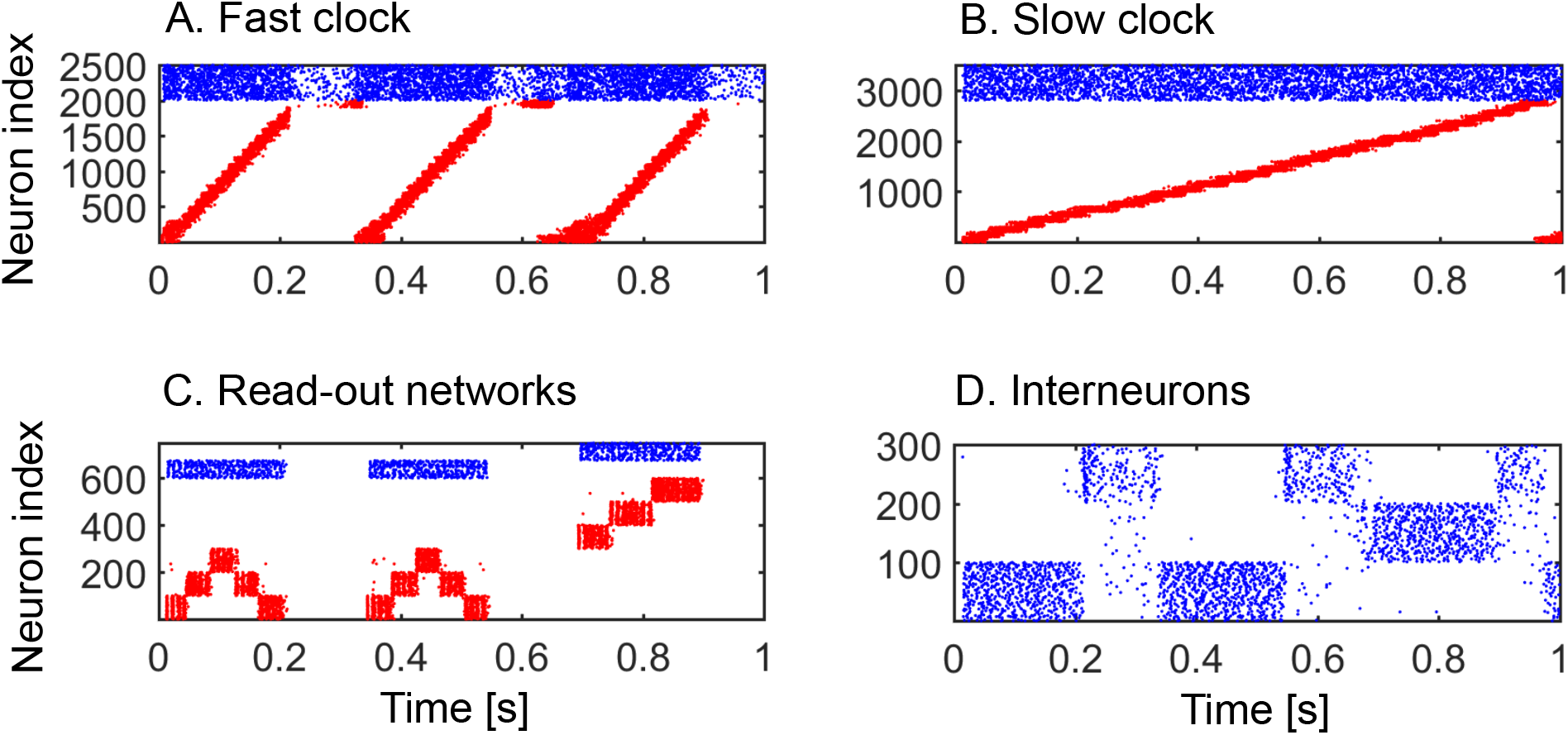
Dynamics of the hierarchical model during target sequence presentation. A. The first cluster of the fast clock receives a high input current at the start of each motif presentation. B. The first cluster of the slow clock receives a high input current at the beginning of the sequence presentation. C. The high input current forces spiking in the read-out neurons. D. The read-out neurons activate the interneurons.

**Fig. S2.**
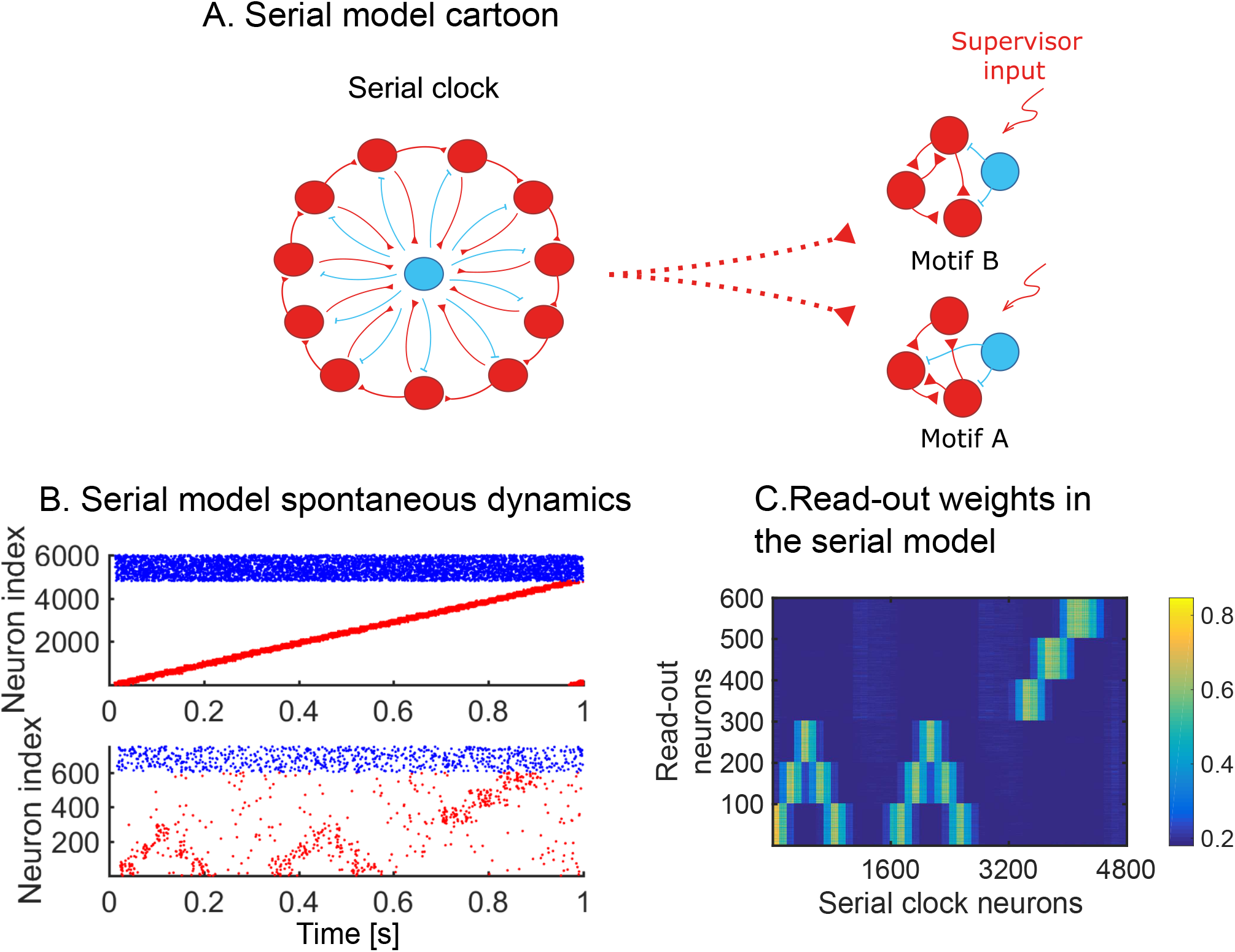
The serial network model. A. A single recurrent network clock (left) produces sequential dynamics and drives the dynamics in the read-out networks (right). The weights from the serial clock to the read-out network are plastic. B. We learn target sequence *AAB*. Spontaneous dynamics is simulated after 90 target sequence presentations. C. The read-out weights after learning. Both motif and syntax information are stored in the same weights.

**Fig. S3.**
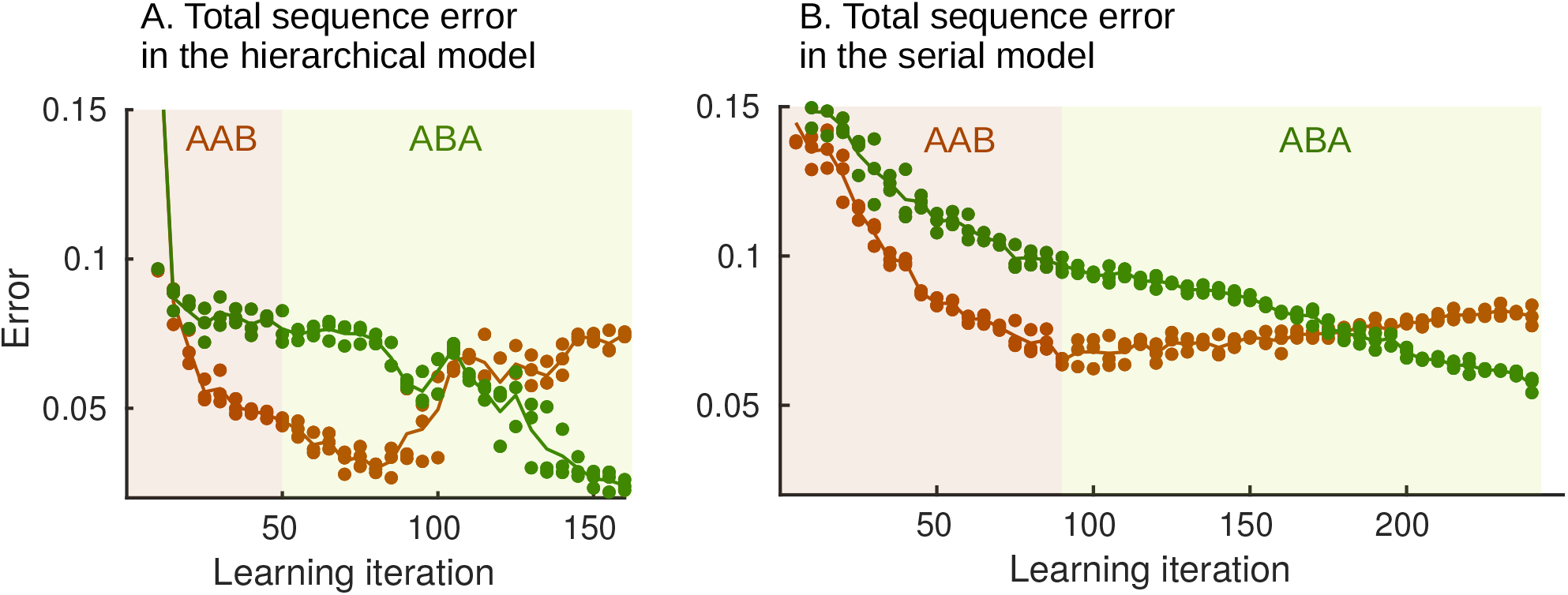
Total sequence error for hierarchical and serial model, during relearning: *AAB* → *ABA*. Spontaneous dynamics is simulated every fifth training iteration and compared with target sequence *AAB* (brown line) and target sequence *ABA* (dark green line) to compute the total sequence error. A. Total sequence error for the hierarchical model. Note how the total sequence error (which is the combination of within-motif error and syntax error) relative to *AAB* decreases for about 30 iterations after target *ABA* is presented for the first time due to the continued improvement in the within-motif dynamics. After this, there is a marked increase in the syntax error and the total error relative to *AAB*. B. Total sequence error for the serial model. The lack of hierarchy in the serial model implies that both the within-motif dynamics and motif ordering has to be relearned. This leads to a more gradual and slower relearning (note the longer x-axis).

**Fig. S4.**
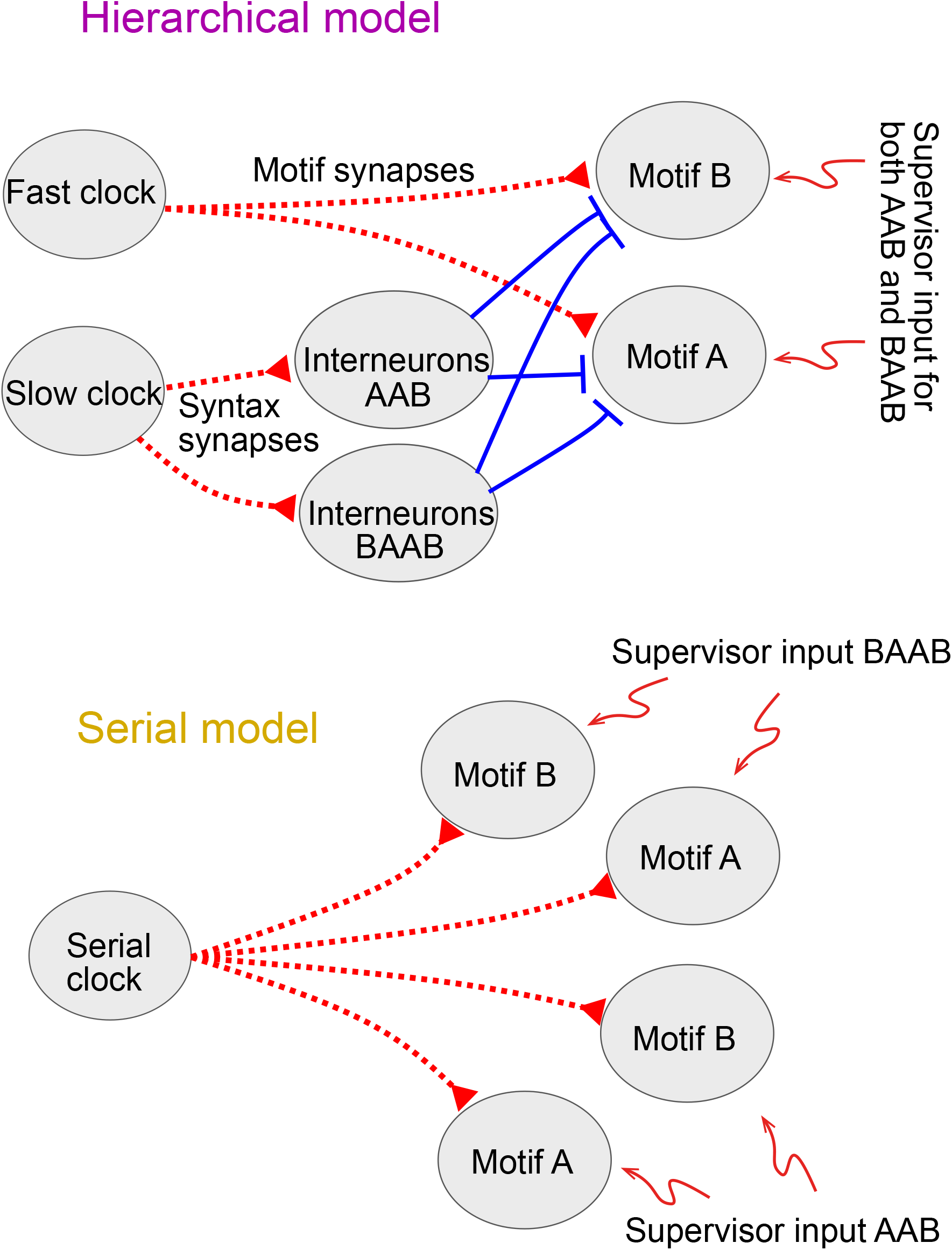
Learning two sequences. The hierarchical model requires an additional interneuron network. An external current is assumed to inhibit the interneurons for sequence *BAAB* when sequence *AAB* is presented and vice versa. The serial model duplicates the entire read-out network. Here also, an external current is assumed to inhibit the read-out networks for sequence *BAAB* when sequence *AAB* is presented and vice versa.

**Fig. S5.**
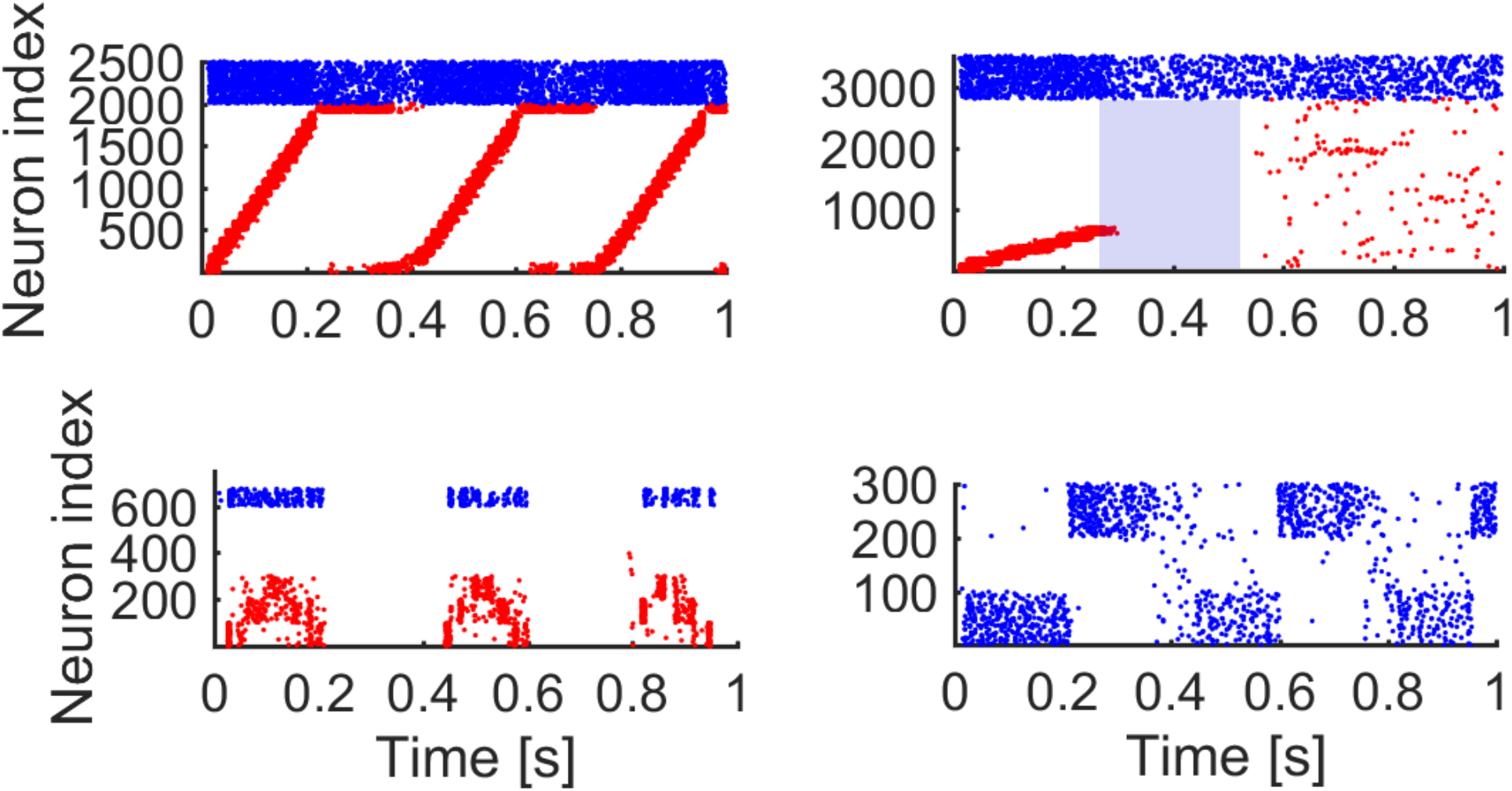
Perturbing the slow clock of the hierarchical network. Blue shade indicates the perturbation time, all excitatory neurons receive no external input for 250 ms. The sequential dynamics in the slow clock breaks down (top right) but random activity in the interneurons (bottom right) leads to sequences in the fast clock (top left), which in turn leads to motif replays (bottom left).

## APPENDIX I. CHANGING THE SLOW CLOCK INTO AN ALL-INHIBITORY NETWORK

The hierarchical model is composed of four networks. These networks can be implemented in various ways. Here, we implement the slow clock differently to illustrate this (Fig. I.1, to be compared with Fig. 1). Sequential dynamics can also be obtained by having an all-inhibitory network (see for example Murray and Escola 2017). Learning the sequence *AAB* with this differently implemented hierarchical model leads to similar results (Fig. I.2, to be compared with Fig. 2). Table I.1 shows the new slow clock inhibitory network parameters. We conserve the other networks. Sequential dynamics in the slow clock is ensured by grouping the inhibitory neurons in 20 clusters of 100 neurons. The inhibitory weights *w*^*II*^ of the same group are multiplied with a factor of 1/30. The inhibitory weights from group *i* to group *i* + 1 mod 20 (*i* = 1..20) are multiplied by a factor of 1/2. This weight structure does not lead to sequential dynamics by itself, some form of adaptation has to be introduced. To this end, short-term depression is used:

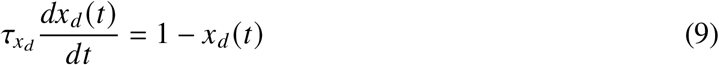

where *x*_*d*_ is a depression variable for each neuron in the all-inhibitory network. This variable is decreased by 0.07 *x*_*d*_ (*t*) when the neuron spikes. The outgoing weights of each neuron in the network are multiplied with this depression variable. The slow clock receives excitatory external random Poisson input and projects to the interneuron networks. The syntax synapses follow the same dynamics as equation 8, but the right hand side of the equation is multiplied by −1 (an inverted STDP window). The parameters are summarized in Table I.2.

**TABLE I.1.** Slow clock inhibitory network parameters

| Constant | Value | Description |
| --- | --- | --- |
| $N^I$ | 2000 | Number of recurrent I neurons |
| $w^{II}$ | 30 pF | I to I synaptic strength |

**Fig. I.1.**
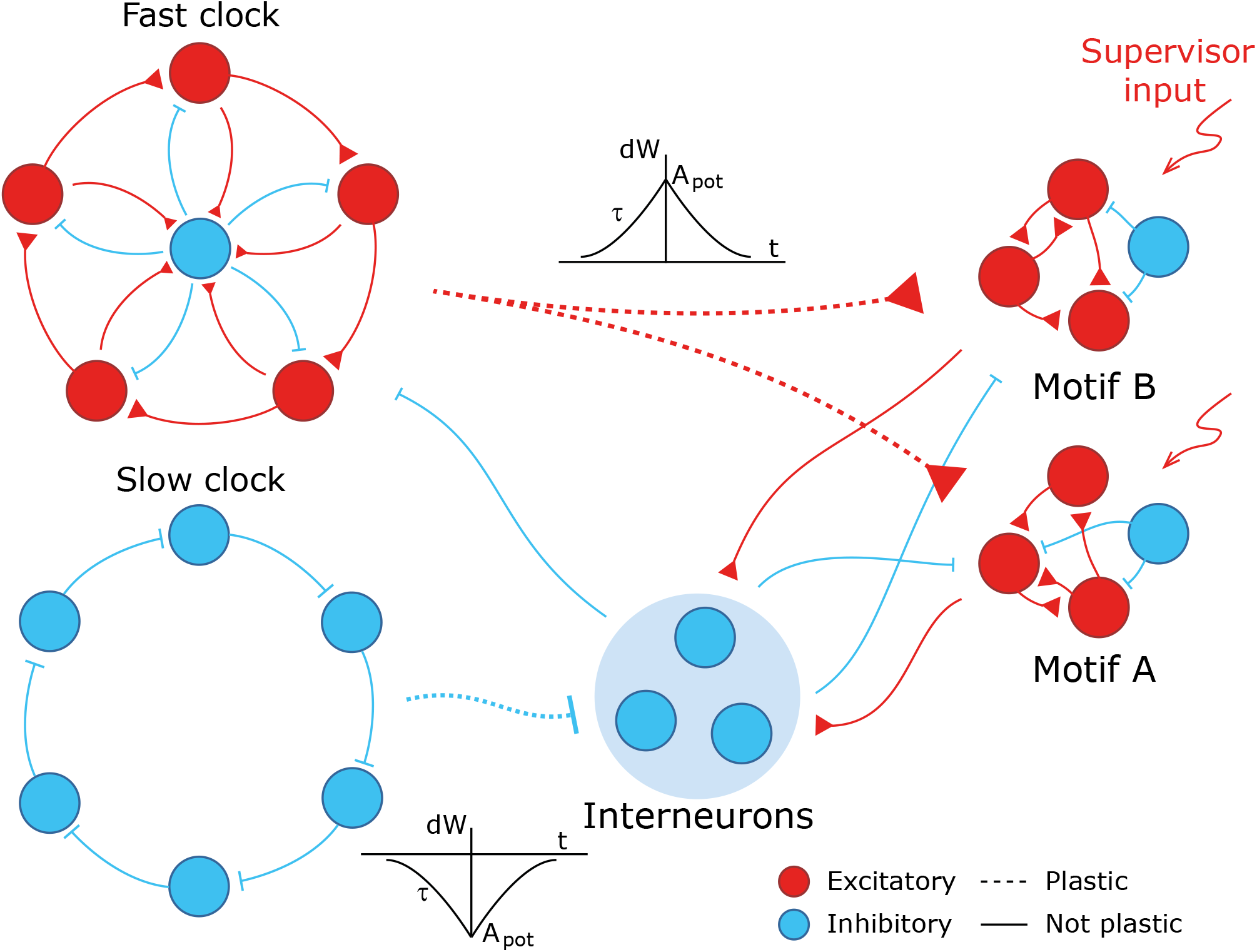
The networks in the model can have different components. Here, the slow clock is replaced by an all-inhibitory network. The syntax synapses follow the same STDP rule as the motif synapses, only inverted.

**TABLE I.2.** Syntax plasticity parameters

| Constant | Value | Description |
| --- | --- | --- |
| $A_{Spot}$ | 0.03 pFHz | Amplitude of potentiation |
| $A_{Sdep}$ | $0.25 \times 10^{-5}$ pFHz | Amplitude of depression |
| $\tau_S$ | 25 ms | Time constant of low pass filter |
| $W_{min}^S$ | 0 pF | Minimum I to I weight |
| $W_{max}^S$ | 0.3 pF | Maximum I to I weight |

**Fig. I.2.**
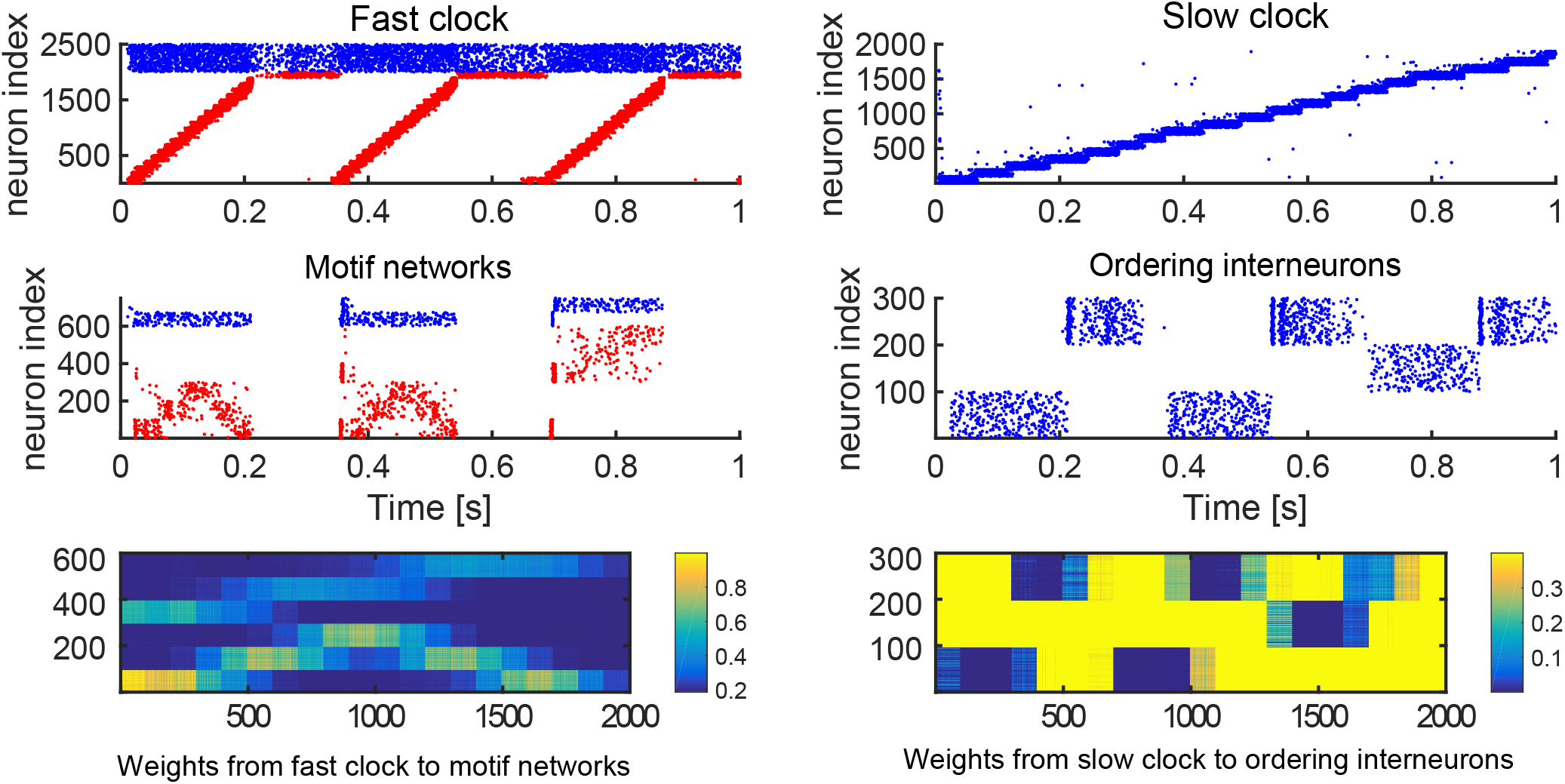
We learn sequence AAB with an inhibitory slow clock network. The simulation shows spontaneous dynamics after training the model for 85 iterations on the sequence *AAB*. The inhibitory neurons of the slow clock inhibit the interneurons now in the correct order (same caption as for Fig. 2).

